# A multi-scale transcriptional regulatory network knowledge base for *Escherichia coli*

**DOI:** 10.1101/2021.04.08.439047

**Authors:** Cameron R. Lamoureux, Katherine T. Decker, Anand V. Sastry, Kevin Rychel, Ye Gao, John Luke McConn, Daniel C. Zielinski, Bernhard O. Palsson

## Abstract

Transcriptomic data is accumulating rapidly; thus, development of scalable methods for extracting knowledge from this data is critical. We assembled a top-down transcriptional regulatory network for *Escherichia coli* from a 1035-sample, single-protocol, high-quality RNA-seq compendium. The compendium contains diverse growth conditions, including: 4 temperatures; 9 media; 39 supplements, including antibiotics; and 76 unique gene knockouts. Using unsupervised machine learning, we extracted 117 regulatory modules that account for 86% of known regulatory network interactions. We also identified two novel regulons. After expanding the compendium with 1675 publicly available samples, we extracted similar modules, highlighting the method’s scalability and stability. We provide workflows to enable analysis of new user data against this knowledge base, and demonstrate its utility for experimental design. This work provides a blueprint for top-down regulatory network elucidation across organisms using existing data, without any prior annotation and using existing data.

**Highlights:**

- Single protocol, high quality RNA-seq dataset contains 1035 samples from *Escherichia coli* covering a wide range of growth conditions
- Machine learning identifies 117 regulatory modules that capture the majority of known regulatory interactions
- Resulting knowledge base combines expression levels and module activities to enable regulon discovery and empower novel experimental design
- Standard workflows provided to enable application of knowledge base to new user data

**Graphical Abstract:** 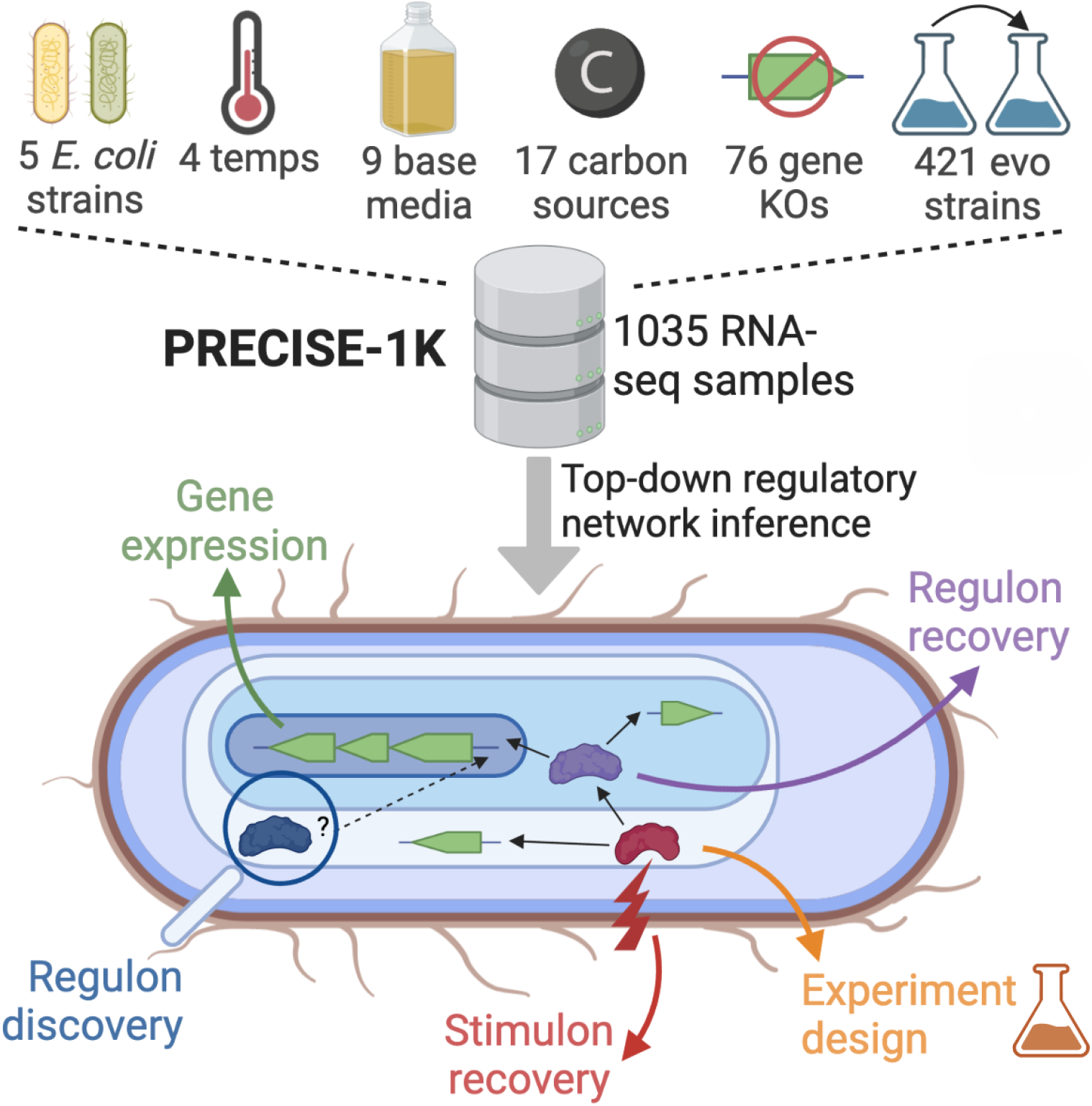

## Introduction

Over the past decade, RNA sequencing (RNA-seq) has emerged as an efficient, high-throughput method to determine the expression state of a cell population. Large RNA-seq datasets (ENCODE Project Consortium, 2012; GTEx Consortium, 2015; Leader et al., 2018; Sastry et al., 2019; Ziemann et al., 2019) have enabled the development and application of machine learning methods to advance our understanding of expression and transcriptional regulation (Avsec et al., 2021; Kelley et al., 2018; Kwon et al., 2020; Sastry et al., 2019; Zhang et al., 2019; Zrimec et al., 2020). As datasets continue to grow, analytic methods must keep pace to convert this data to biological knowledge. In particular, top-down elucidation of a transcriptional regulatory network (TRN) directly from RNA-seq data would address a pressing need.

Independent component analysis (ICA) (Comon, 1994) is a signal processing algorithm that outperforms other methods for the extraction of biologically meaningful regulatory modules from gene expression data (Saelens et al., 2018). Application of this method to publicly-available prokaryotic expression data has consistently recovered TRN modules across organisms (Chauhan et al., 2021; Poudel et al., 2020; Rajput et al., 2022; Rychel et al., 2020; Sastry et al., 2019; Yoo et al., 2022). ICA’s effectiveness results from its ability to identify independent groups of genes that vary consistently across samples, regardless of group size or overlapping membership. As a result, a complete top-down TRN extraction is dependent upon an input dataset with sufficient scale and diversity in conditions to activate a broad range of regulatory signals.

Large RNA-seq datasets compiled from multiple sources can be subject to batch effects that confound analysis. Mitigating these effects remains an important goal and an active area of research (Liu and Markatou; Zhang et al., 2020). Single-protocol, high-quality, curated RNA-seq datasets provide an alternative to batch effect correction by obviating them entirely. However, amassing the quantity of data necessary to perform systems-level inference is a challenge.

To simultaneously address these issues, we assembled PRECISE-1K, a 1035-sample, single-protocol RNA-seq dataset for the key model organism *Escherichia coli* K-12 MG1655. The thousand-sample-scale *<u>P</u>*recision *<u>E</u>*NA-seq *<u>E</u>*xpression *<u>C</u>*ompendium for *<u>I</u>*ndependent *<u>S</u>*ignal *<u>E</u>*xtraction contains 38% of all publicly-available high-quality RNA-seq data for *E. coli* K-12 and includes a broad range of growth conditions. Here, we present a multi-scale analysis of PRECISE-1K. Specifically, we: (1) describe genome-wide gene expression patterns across conditions; (2) use ICA to extract a top-down TRN comprising 117 *i*ndependently *modulated* groups of genes (iModulons); (3) describe systems-level transcriptome properties; (4) discover novel regulons for two putative transcription factors; (5) propose a promoter sequence basis for two regulatory modules; (6) add 1675 high-quality publicly available K-12 samples to PRECISE-1K and extract similar regulatory modules; and (7) demonstrate a workflow for systems-level transcriptome analysis of external data using our top-down TRN. This example workflow, along with all analyses presented here, are available for use at our GitHub repository, https://github.com/SBRG/precise1k. Our TRN, along with those for the other organisms mentioned above, can be explored at iModulonDB.org (Rychel et al., 2021). The PRECISE-1K compendium and the top-down TRN derived from it comprise a multi-scale transcriptomic knowledge base. The analyses enabled by this knowledge base highlight its potential to play a key role in multi-omic systems-level investigations of this critical model organism for cellular biology, pathogenicity, and synthetic biology. Moreover, our knowledge base provides a useful resource for informing novel experimental designs. Ultimately, PRECISE-1K and iModulons combine to provide a blueprint for top-down TRN extraction across organisms without dependence on prior annotation.

## Results

### PRECISE-1K is a large, single-protocol, and high-quality RNA-seq compendium

We constructed PRECISE-1K to enable a multi-scale analysis of the *E. coli* K-12 MG1655 TRN (**Supplemental Figure 1**). PRECISE-1K is a large, high-fidelity expression compendium consisting of 1035 RNA-seq samples generated by a single research group using a standardized experimental and data processing protocol (see **Methods**). The samples come from 45 distinct projects. PRECISE-1K constitutes a nearly 4-fold increase in size from the 278-sample PRECISE (Sastry et al., 2019) (**Figure 1A**). Replicates are tightly correlated, with a median Pearson’s *r* of 0.99 (**Figure 1C**).

**Figure 1:**
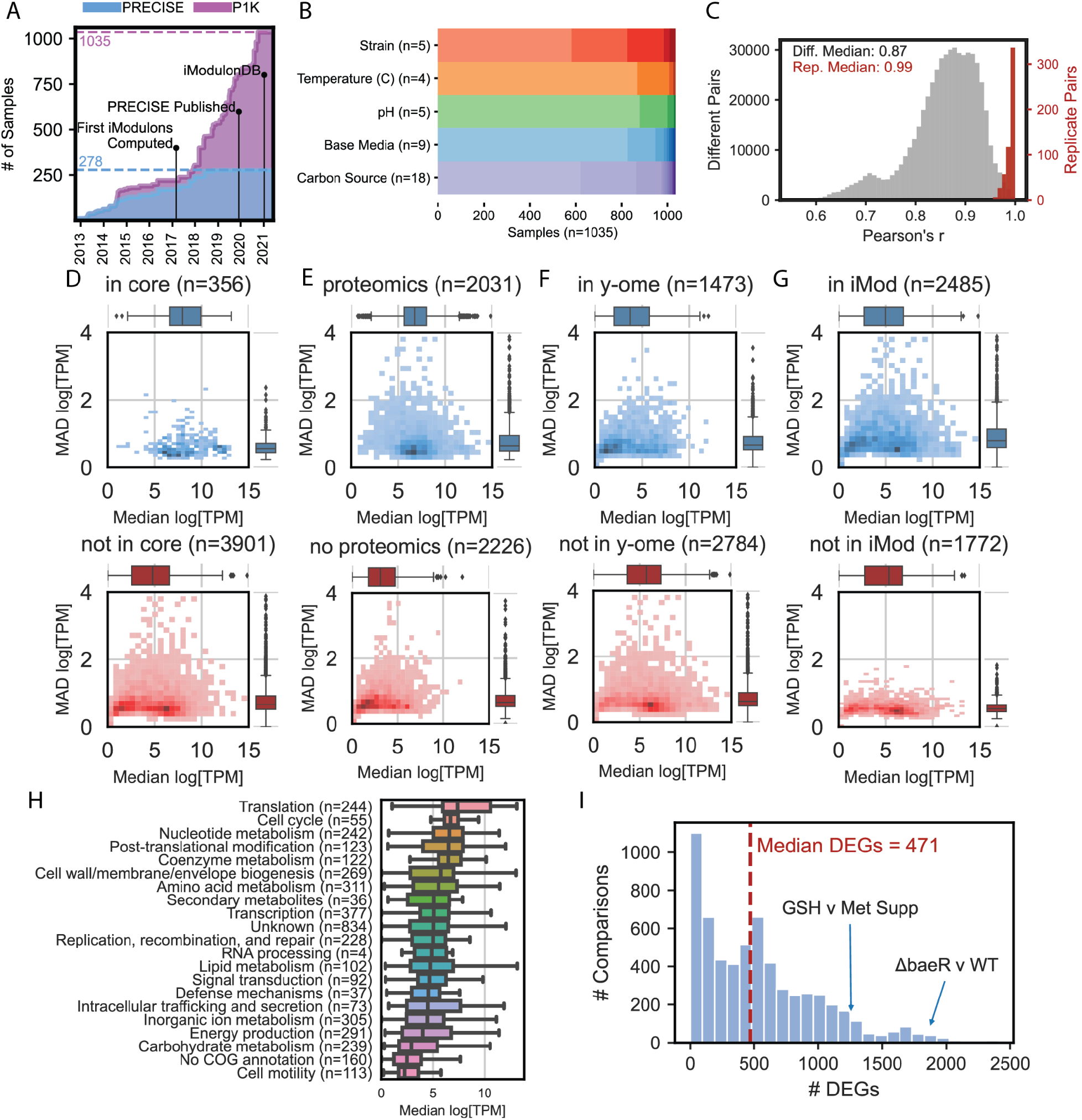
PRECISE-1K reveals expression trends in the *E. coli* transcriptome. **A)** The growth in single-protocol transcriptomics samples contained in the PRECISE to PRECISE-1K databases. **B)** Summary of selected conditions for PRECISE-1K samples. For a full breakdown, see Supplemental Figure 2. **C)** Histogram of Pearson’s *r* for both all replicate pairs and all non-replicate pairs. Samples included in PRECISE-1K are required to have replicate correlations of at least 0.95. **D-G)** Breakdown of gene expression and expression variance by category. in core = core *E. coli* genome based on pangenomic analysis (Norsigian et al., 2020); proteomics = proteomics data available (Heckmann et al., 2018; Schmidt et al., 2016); y-ome = poorly-annotated genes (Ghatak et al., 2019); in iMod = appears in at least 1 iModulon. **H)** Breakdown of gene expression by COG category across PRECISE-1K. **I)** Histogram of the number of differentially expressed genes (DEGs) computed between condition pairs within the same project. GSH = glutathione, Met = methionine.

PRECISE-1K comprises a wide range of growth conditions, including: 5 strains; 4 temperatures, 5 pHs, 9 base media, 18 carbon sources, 38 supplements, 76 unique gene knockouts, 421 evolved samples, and 87 fed-batch cultures (**Figure 1B**). PRECISE-1K features projects involving: adaptation to new growth conditions (Anand et al., 2019, 2020; Chen et al., 2021; Du et al., 2020; McCloskey et al., 2018); expression of heterologous (Tan et al., 2020) and orthologous (Sandberg et al., 2020) genes; and a genome-reduced strain (Hirokawa et al., 2013). PRECISE-1K thus represents a broad range of conditions under which changes in the composition of the *E. coli* transcriptome may be studied.

Principal component analysis (PCA) of PRECISE-1K reveals minimal batch effects. The first two principal components of the dataset capture 26.8% of the overall variance (**Supplemental Figure 3**). The samples do not cluster notably by the RNA-seq library preparer, indicating an absence of a commonly observed batch effect (Zhang et al., 2020). Any larger separation between samples in principal component space is explained by differences between project growth conditions. Projects that use diverse growth media (e.g. two-component system project (Choudhary et al., 2020) and antibiotic/media project (Sastry et al., 2020)) and projects that significantly alter the genome (minicoli, a genome-reduced *E. coli* strain) account for sample distinction in principal component space.

### PRECISE-1K highlights global and environment-specific gene expression patterns

PRECISE-1K provides a high-level view of absolute expression and expression variance across the *E. coli* genome. Interrogating these systems-level expression patterns reveals the hallmarks of the transcriptome.

Genes in the core genome (i.e. genes shared across all *E. coli* strains, as defined by pangenomic analysis (Norsigian et al., 2020)) are significantly more expressed than non-core genes (*P*=2.6E-105, Mann-Whitney U test, *m*=356, *n*=3901) (**Figure 1D**). The core genome also exhibits significantly less variation in expression (*P*=1.0E-15). 43% (152/356) of core genome genes are in the clusters of orthologous genes (COG) category “Translation, ribosomal structure and biogenesis,” accounting for 62% (152/244) of all genes in that COG category. Indeed, this COG category has the highest median expression across PRECISE-1K (**Figure 1H**). Taken together, these results underscore the importance of maintaining consistently high expression levels for these genes across a wide range of growth conditions.

Additionally, genes for which proteomics data is available (Heckmann et al., 2020; Schmidt et al., 2016) have significantly higher expression (*P*=1.2E-150, *m*=2031, *n*=2226), consistent with a known bias towards higher-expressed genes amongst proteomics samples (**Figure 1E**). We also compared the expression of poorly-annotated genes (referred to as the “y-ome” in *E. coli* (Ghatak et al., 2019)) to genes with more complete annotation. y-ome genes have significantly lower expression (*P*=1.0E-75, *m*=1473, *n*=2784) than non-y-genes, highlighting the lack of transcription in standard laboratory conditions as a potential reason for these genes’ relative lack of annotation (**Figure 1F**).

We performed differential gene expression analysis within each member project for all projects in the PRECISE-1K compendium. A median of 471 differentially expressed genes (DEGs) were found across all pairwise within-project comparisons (**Figure 1I**). Many comparisons produced close to 0 DEGs - for example, comparison of a *qseF* deletion to a wild-type control after six hours of batch culture yielded only six DEGs. Other in-project comparisons yielded far more DEGs. For example, the comparison between wild-type growth in minimal media and deletion of two-component system (TCS) response regulator *baeR* with ethanol supplementation yielded 1868 DEGs.

### Top-down regulatory modules capture transcriptome allocation and the transcriptional regulatory network

We used the signal extraction machine learning algorithm independent component analysis (ICA) to identify 201 independently modulated groups of genes (iModulons) from PRECISE-1K. iModulons are quantitative representations of regulatory modules that contain a regulator’s target genes and quantify its activity level in each sample. The 201 iModulons extracted from PRECISE-1K reconstruct 83% of the total variance in the dataset. 117 of these iModulons are classified as Regulatory, as they are significantly enriched in genes belonging to a known regulon (**Figure 2A;** see **Supplemental Data** for a full summary of all 201 iModulons including regulator enrichment statistics). These Regulatory iModulons explain 56% of the total variance in PRECISE-1K. 36 genomic iModulons that capture known genetic alterations (e.g. gene knockouts) and 17 biological iModulons (comprised of genes with shared function but lacking significant regulon enrichment) account for another 19% of the variance. 9 uncharacterized iModulons account for just 6% of the variance in the dataset. Altogether, 94% of the variance captured by iModulons can be explained by either regulatory, genomic, or biological phenomena.

**Figure 2:**
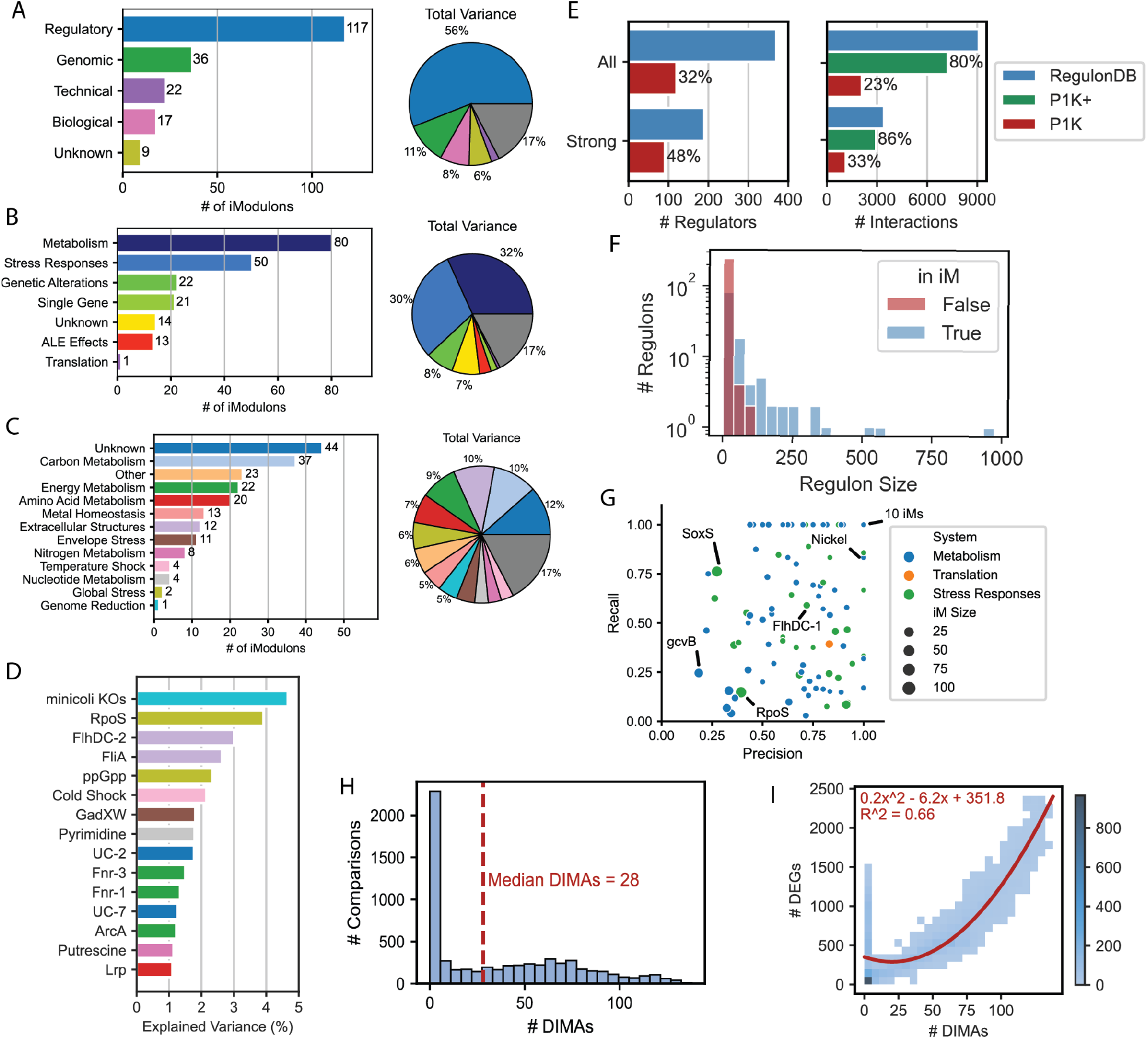
PRECISE-1 K has 201 iModulons that represent a range of cellular processes. **A)** A breakdown of PRECISE-1K iModulons by their annotation category: ‘Regulatory’ denotes significant enrichment of one or more known regulators; ‘Technical’ includes a single gene or technical artifact iModulon; ‘Genomic’ includes iModulons related to known genomic interventions (i.e., knockouts or segmental amplifications due to adaptive laboratory evolution); and ‘Biological’ includes iModulons containing genes of related function without significant regulator enrichment, or pointing to potential new regulons. Pie chart denotes iModulon annotation categories by percentage of variance explained. Gray wedge indicates variance unexplained by iModulons. **B)** Breakdown of iModulons based on system annotation. ALE = adaptive laboratory evolution. **C)** Breakdown of iModulons based on functional annotation. **D)** Top 15 iModulons ranked by % of variance explained. Color code matches panel **C**. **E)** Comparison of regulators and regulatory interactions recovered by PRECISE-1K and available in RegulonDB. All = all evidence levels; Strong = only strong evidence interactions per RegulonDB; P1K+ = all interactions for which the corresponding regulator is captured by PRECISE-1K. **F)** Histogram of regulon size for regulators captured or not captured by at least one iModulon. **G)** Summary of precision and recall for 117 regulatory iModulons with RegulonDB regulons as reference. **H)** Histogram of differential iModulon activities (DIMAs) for same condition comparisons as Figure 2I. **I)** Comparison of number of DEGs and DIMAs for the same condition pairs. The best fit curve is shown in red.

iModulons extracted from PRECISE-1K reconstruct a significant fraction of the total regulatory interactions available in RegulonDB (Santos-Zavaleta et al., 2019), the premier database for curated and validated regulatory network information for *E. coli.* 32% of all known regulatory molecules (and 48% with strong evidence) are captured by regulatory iModulons (**Figure 2E**). Moreover, 23% of all specific regulatory interactions (33% of strong-evidence interactions) are reconstituted in iModulons. iModulons are known to capture regulatory signals by identifying the most strongly-regulated genes in a regulon based on promoter sequence (Qiu et al., 2022). This likely accounts for the relatively lower precision and recall enrichment statistics observed for larger iModulons that capture more global regulators (**Figure 2G**). Thus, considering a regulatory iModulon as a biomarker for all of its regulator’s interactions reveals that iModulons in fact reconstitute 80% of all known regulatory interactions (86% when considering only strong evidence). Importantly, iModulons preferentially capture the signals of larger regulons (**Figure 2F**), increasing their utility in describing transcriptome state across growth conditions.

58% of genes (2485/4257) are members of at least one iModulon. These genes have higher expression variation than genes not present in any iModulons (*P*=1.2E-219, Mann-Whitney *U* test, *m*=2485, *n*=1772). However, median expression itself does not depend significantly on membership (*P=*0.28) (**Figure 1G**). Thus, iModulon membership is not restricted to higher-expressed genes. Indeed, 56% (823/1473) of y-ome genes - demonstrated above to be significantly less expressed - are members of at least one iModulon, highlighting the potential for iModulons to uncover putative functions for these uncharacterized genes. The median iModulon consists of 10 genes, though many iModulons are much larger, such as global stress responses RpoS (122 genes) and SoxS (117) (**Supplemental Figure 4A**). 35% of genes in an iModulon (879/2485) are members of 2 or more iModulons, with 2 genes *(ynfM* and *bhsA)* appearing in 7 each (**Supplemental Figure 4B**). These characteristics highlight iModulons’ ability to capture overlapping regulatory modules of varying scale.

80 metabolism and 50 stress response iModulons account for 32% and 30% of the variance in PRECISE-1K, respectively (**Figure 2B**). This breakdown reveals that, in aggregate across PRECISE-1K growth conditions, *E. coli* allocates transcriptome nearly equally to these two critical functions, emphasizing a “fear-greed” tradeoff. Interestingly, the numbers of iModulons for these two functions differ considerably; the cell thus has a tendency towards more diversified regulation for metabolic capabilities and more centralized control for stress responses. Indeed, just two iModulons - RpoS and ppGpp, major stress response regulators - account for 6% of the variance in the dataset (**Figure 2C**).

iModulons capturing the signals of global regulators (regulators with more than 25 regulatory targets) account for large proportions of the overall variance in the dataset. Flagella-related regulators FlhDC and FliA in combination explain over 5% of the expression variance, while anaerobic growth regulators FNR and ArcA combine to explain over 3% of the variance (**Figure 2D**). These insights highlight the ability of global regulators to mobilize large-scale transcriptomic responses. Indeed, these regulators (along with iron regulator Fur) are responsible for variance between wild-type control samples run across projects, despite overall tight correlation between those samples (**Supplemental Figure 5**).

### Regulatory modules enable systems-level analysis of transcriptome states

Because iModulons include an explicit representation of regulator activity levels, they enable differential iModulon activity (DIMA) analysis. DIMA analysis allows for a systems-level comparison of transcriptome states by reducing hundreds or thousands of DEGs to a median of just 28 iModulons (**Figure 2H**). On average, a comparison between any two conditions in PRECISE-1K yields almost twenty times fewer differentially-activated iModulons than DEGs (**Figure 2I**). DEGs scale quadratically with DIMAs, highlighting the particular usefulness of DIMA analysis for systems-level transcriptional analysis.

iModulon activities reflect the overall activity state of a transcriptional regulator across environmental conditions in PRECISE-1K. A stimulon is a higher-level regulatory network composed of multiple regulons that respond to a particular stimulus (**Supplemental Figure 1**). While iModulons, by definition, include independently modulated groups of genes, in many instances these independent groups of genes are regulated in response to similar environmental stimuli, thus forming a stimulon. Two-component systems (TCS) - composed of a membrane-bound sensor and a cytoplasmic response regulator - enable the cell to sense and respond to important extracellular signals. iModulons derived from PRECISE-1K capture the response signal for 15 of 27 known TCS response regulators, providing insight into the cell’s regulatory response to critical stimuli such as nitrogen, inorganic phosphate, and alkali metals.

Additionally, iModulons can be clustered based on their activities to reveal stimulons. For example, one cluster captures the joint regulation of flagella formation by transcription factor complex FlhDC and sigma factor FliA (σ^28^) (**Supplemental Figure 6**). Six iron-related iModulons, five anaerobiosis-related iModulons, and four amino acid-related iModulons also group together in this activity-based fashion. Thus, iModulons in combination can shed light on broad transcriptome patterns and the hierarchy in the transcriptional regulatory network.

### PRECISE-1K enables regulon discovery

Functional annotation for putative TFs remains elusive (Gao et al., 2018). However, iModulons are a powerful tool for the discovery and analysis of new regulons. PRECISE elucidated the regulons for three previously uncharacterized TFs (YieP, YiaJ/PlaR, and YdhB/AdnB), and expanded the regulons of three known TFs (MetJ, CysB, and KdgR) (Sastry et al., 2019). Many of these regulatory interactions were confirmed through DNA-binding profiles (Rodionova et al., 2020a, 2020b; Sastry et al., 2019). Furthermore, three novel regulons were predicted from iModulons derived from a microarray dataset (Sastry et al., 2021a). iModulons from PRECISE-1K recapitulate these previous results and reveal two new potential regulons.

The putative YgeV regulon contains 13 genes, of which 7 are putatively involved in nucleotide transport and metabolism (**Figure 3A**). YgeV is predicted to be a Sigma54-dependent transcriptional regulator, and Sigma54-dependent promoters were previously identified upstream of the *xdhABC* and *ygeWXY* operons, which are in the YgeV iModulon (Reitzer and Schneider, 2001). Although the iModulon does not contain the gene *ygeV, ygeV* is divergently transcribed from *ygeWXY.* A prior study (DeLisa et al., 2001) found that expression of *ygfT* was reduced in a YgeV mutant strain. Since *ygfT* is in the YgeV iModulon, this indicates that YgeV may serve as an activator for the genes in its iModulon. The activity of the YgeV iModulon rarely deviates from the reference condition; however, it is most active when knockouts of TCS response regulators BaeR or CpxR are exposed to ethanol (**Figure 3B**). Therefore, we predict that the TF YgeV responds (either directly or indirectly) to ethanol to activate genes related to purine catabolism, and is repressed by TCS BaeRS and CpxAR.

**Figure 3:**
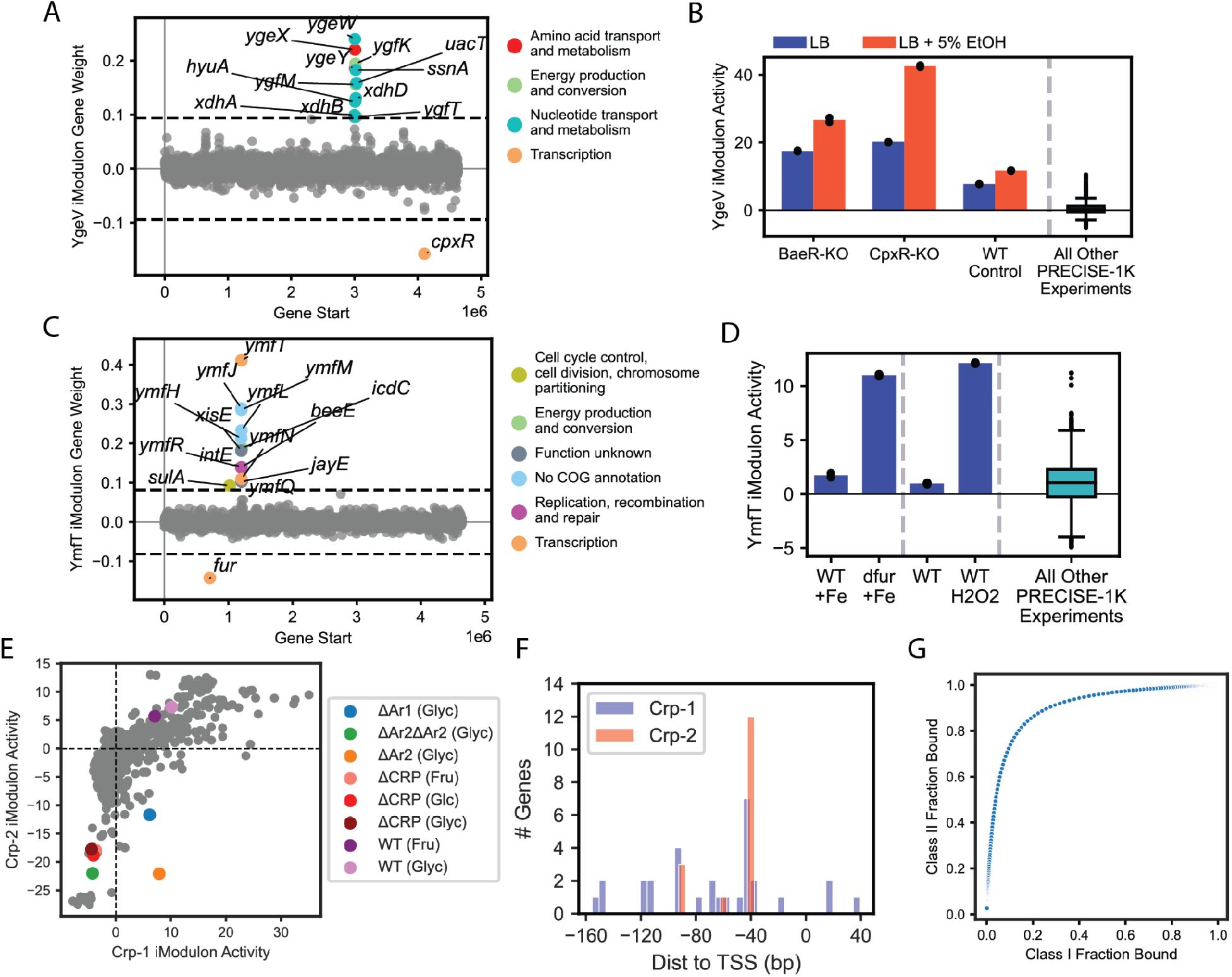
iModulons predict and stratify regulons. **A)** iModulon gene weights for the putative YgeV iModulon vs. genome position. **B)** Activity of the YgeV iModulon in different media conditions. Each colored bar is the mean of two biological replicates (shown as individual black points). **C)** iModulon gene weights for the putative YmfT iModulon vs. genome position. **D)** Activity of the YmfT iModulon in different media conditions. Each colored bar is the mean of two biological replicates (shown as individual black points). **E)** Activity tradeoff between Crp-1 and Crp-2 iModulons. Colored points from samples involving partial and total CRP deletions. **F)** Histogram of CRP binding site locations for Crp-1 and Crp-2 iModulons. TSS = transcription start site of transcription unit for each gene. Data from RegulonDB. **G)** Simulated binding curve for CRP Class I and Class II promoters. Each point indicates a particular CRP concentration. Binding modeled as 10x tighter at Class II vs Class I promoters.

The putative YmfT regulon contains 15 genes, including *ymfT* itself. It contains 12 of the 23 genes in the e14 prophage (Mehta et al., 2004) (**Figure 3C**). The putative YmfT iModulon is most active in strains lacking the ferric uptake regulator Fur, or in strains challenged by oxidative stress through hydrogen peroxide (**Figure 3D**). Absence of Fur leads to overproduction of iron uptake proteins, oxidative damage, and, subsequently, mutagenesis (Touati et al., 1995). Therefore, we predict that YmfT responds to oxidative stress to alter the expression of the e14 prophage.

These examples illustrate the potential for iModulons to predict new regulons and identify optimal conditions to study their activities. **Supplemental Table 2** lists all regulons that have been identified or modified using PRECISE-1K and its associated iModulons.

### iModulons capture promoter-level mechanisms of Crp regulation

ICA discovers independent sub-groups of genes within global regulons that exhibit distinct regulatory dynamics. For example, previous work has shown how the Fur-1 and Fur-2 iModulon activities (both of which are in the Fur regulon) vary with iron availability (Sastry et al., 2020). Here, we demonstrate that iModulons reflect biochemical mechanisms of TF binding by examining the relationship between two iModulons - Crp-1 and Crp-2 - that capture parts of the CRP regulon. CRP contains multiple RNA polymerase-interacting domains (Ar1-3) (Lawson et al., 2004) that facilitate its binding to Class I and Class II promoters. Class I promoters canonically involve binding centered 61.5 base pairs upstream of the transcription start site, and Class II are centered 41.5 base pairs upstream (Busby and Ebright, 1999).

The activities of the Crp-1 and Crp-2 iModulons across all PRECISE-1K conditions form a distinct nonlinear relationship (**Figure 3E**). As expected, low activities of both iModulons correspond with deletion of CRP, which is known to activate most of the genes in the two iModulons. Deletion of the Ar2 binding domain - implicated in Class II regulation - results in some Crp-1 activity but no Crp-2 activity (orange dot in **Figure 3E**). CRP binding sites for genes unique to Crp-1 are broadly distributed around the canonical Class I binding location, while Crp-2-specific genes have CRP binding sites more consistently at the Class II location (**Figure 3F**). A steady-state biophysical model with 10-fold different binding affinities for Class I and Class II binding sites yields a similar binding site occupancy relationship as that between the Crp iModulon activities (**Figure 3G**). From this evidence, we propose that the Crp-1 and Crp-2 iModulons correspond to Crp regulatory activity at Class I and Class II promoter genes, respectively. This analysis highlights the capability of PRECISE-1K iModulons to capture regulatory dynamics within a single regulon.

### Expanded dataset highlights regulatory modules’ robustness and PRECISE-1K’s quality

To further expand our dataset, we sourced all publicly-available RNA-seq data for *E. coli* strain K-12 from NCBI’s Sequence Read Archive (SRA) (Leinonen et al., 2011). From 3,230 K-12 samples, our processing and quality control pipeline yielded 1,675 high-quality K-12 expression profiles. In combination with PRECISE-1K, these data created the K-12 dataset, a high-quality transcriptomics dataset consisting of 2,710 expression profiles (**Figure 4A**). These profiles come from 134 different projects, including 15 K-12 substrains and 9 distinct temperatures and pHs (**Figure 4B**). ICA decomposition of the K-12 dataset yields 194 iModulons.

**Figure 4:**
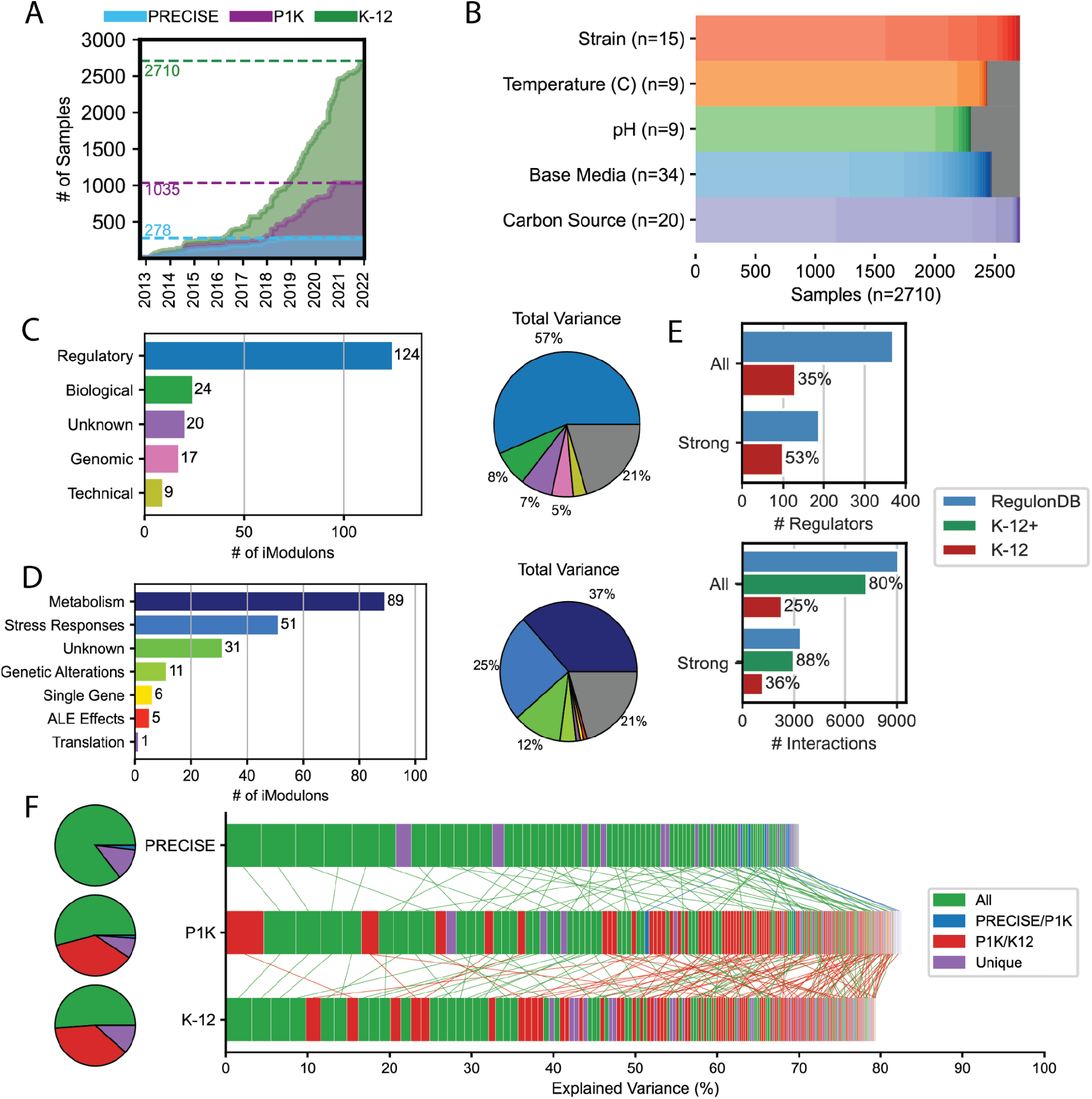
Adding public K-12 data to PRECISE-1K highlights PRECISE-1K’s stability. K-12 is a combined dataset composed of PRECISE-1K (1035 samples) plus all publicly-available high-quality RNA-seq data for *E. coli* K-12 (1675 samples). **A)** The accumulation of high-quality RNA-seq data for K-12 over time. **B)** Summary of selected conditions for K-12 samples. Gray indicates samples for which the public metadata did not specify the relevant variable. **C)** K-12 iModulons by their annotation category (see **Figure 2A** legend). Pie chart denotes iModulon annotation categories by percentage of variance explained. The 194 annotated iModulons together explain 81% of the variance. Gray wedge indicates variance unexplained by iModulons. **D)** Same as **C** but based on specific system annotation. **E)** Comparison of regulators and regulatory interactions recovered by K-12 and available in RegulonDB. All = all evidence levels; Strong = only strong evidence interactions per RegulonDB; K-12+ = all interactions for which the corresponding regulator is captured by the K-12 dataset. **F)** Comparison of iModulons from three RNA-seq datasets: PRECISE (Sastry et al., 2019); PRECISE-1K (this paper); and K-12. Explained variance is within each dataset. Each small rectangle represents an iModulon for the corresponding dataset, ordered left to right in descending order of explained variance. Lines link iModulons with Spearman’s correlation coefficient greater than 0.25. Green = correlated set of iModulons exists across all 3 datasets; blue = correlated set of iModulons only exists in PRECISE/PRECISE-1K; red = correlated set for PRECISE-1K/K-12 only; gray = iModulon unique to dataset. Pie charts indicate fraction of dataset variance explained in each correlation category.

The distribution of iModulons by category – both in number and by explained variance – is broadly similar to that of PRECISE-1K. Regulatory iModulons account for 64% of the total number, and 57% of the total variance in the dataset (**Figure 4C**). Percentages of variance explained by metabolic (37%) and stress (25%) iModulons indicate that the K-12 dataset is relatively biased towards metabolic changes compared with PRECISE-1K (**Figure 4D**). Coverage of known regulatory network interactions increases only minutely as compared with PRECISE-1K alone, despite the more than doubling of the dataset’s size (**Figure 4E**). Indeed, 89% of K-12’s explained variance comes from iModulons already identified in either PRECISE or PRECISE-1K. In contrast, 45% of explained variance from PRECISE-1K comes from iModulons not present in PRECISE (**Figure 4F**). iModulons can also explain a slightly larger fraction of variance in PRECISE-1K than in the K-12 dataset. Taken together, these results suggest that PRECISE-1K has sufficient scale and condition variety to represent the *E. coli* TRN, and further large-scale additions of data may provide diminishing returns.

### Applying the PRECISE-1K knowledge base to new data: a case study

PRECISE-1K and its associated suite of analyses are designed to be applicable to new *E. coli* RNA-seq datasets. We demonstrate this capability for one project from the public K-12 dataset. This project - called AAT for anaerobic-aerobic transition - captured six time-points in triplicate from 0 to 10 minutes after aeration of a previously anaerobic chemostat culture of *E. coli* K-12 W3110 (Bui and Selvarajoo, 2020). PRECISE-1K iModulon activities for the AAT project can be inferred without necessitating full re-processing through the entire workflow. These inferred activities in turn enable analysis of AAT’s samples both within the project and within the context of all PRECISE-1K’s samples. The Jupyter notebook used for this case study is available at https://github.com/SBRG/precise1k and is set up to be used for analysis of any new data.

We identified the iModulons with unusually divergent activities in AAT compared to the rest of PRECISE-1K (**Figure 5A**). iModulons related to energy metabolism featured prominently; for example, the formate hydrogen lyase (FHL) iModulon had a maximum absolute activity in AAT six standard deviations off the PRECISE-1K average. FHL is known to be active under anaerobiosis during glucose fermentation. An activity histogram further contextualizes these observations: while aerobic metabolism regulator ArcA is over three standard deviations away from the PRECISE-1K average at maximum in AAT, other AAT samples are closer to the PRECISE-1K median (**Figure 5B**). To further characterize iModulon activity changes within AAT, DIMA analysis can identify iModulons that change significantly between any two sets of samples. Comparing aeration onset to 10 minutes post-aeration highlights the roles of key energy metabolism global regulators in facilitating this transition (**Figure 5C**). Anaerobic metabolism global regulator Fnr is more active at onset, while aerobic metabolism regulator ArcA and global iron regulator Fur increase in activity 10 minutes after aeration. Fnr’s activity decreases nonlinearly following aeration of the culture, reducing its reference level in aerobic culture within 5 minutes (**Figure 5D**). Activity clustering highlights increased activity of the anaerobic stimulon at aeration onset, followed by increased activation of the iron stimulon 10 minutes post-aeration (**Supplemental Figure 7**).

**Figure 5:**
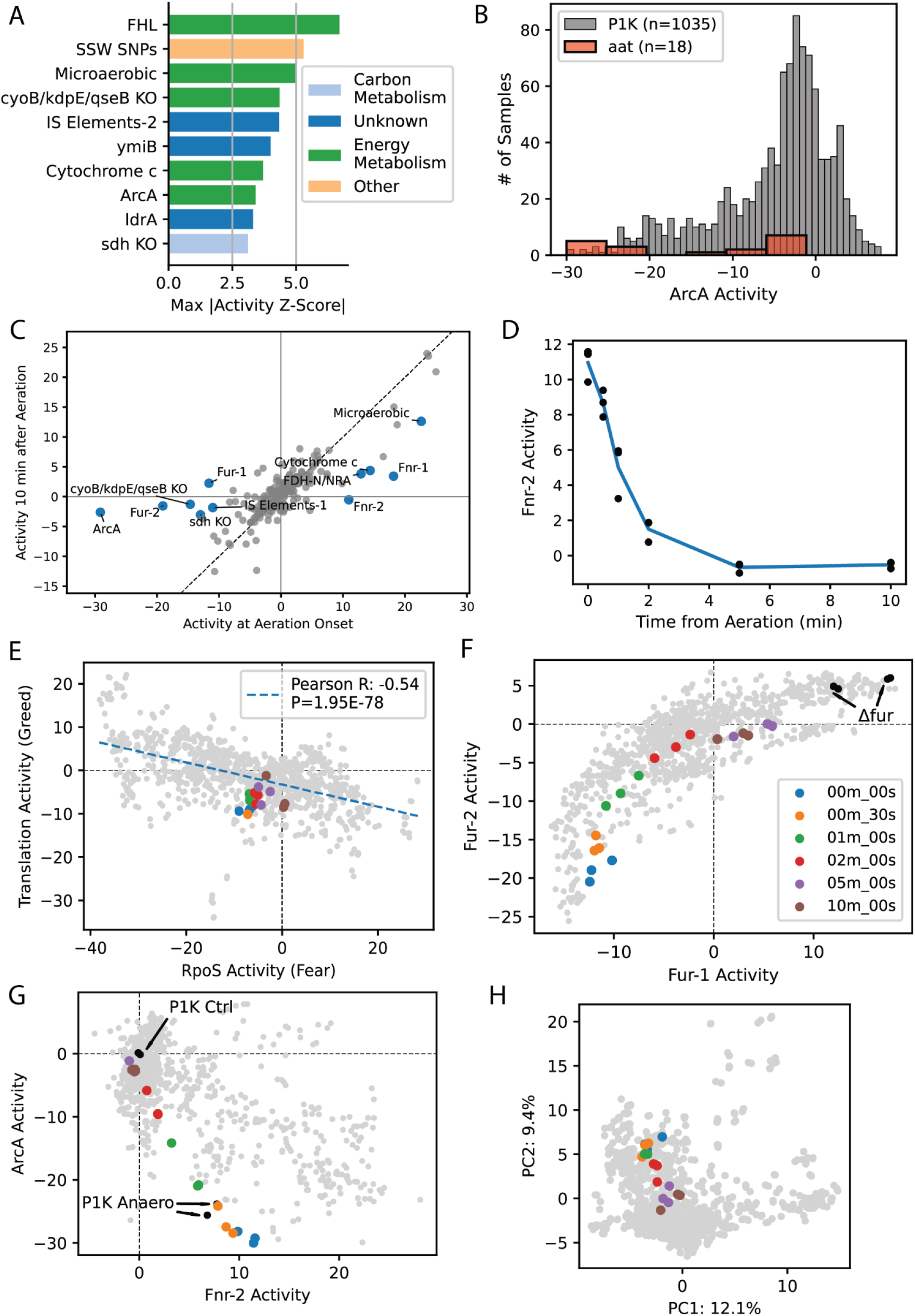
PRECISE-1K and iModulons provide key insight for assessing systems-level transcriptome changes for new data. For all panels in this figure, the example new data comes from the public K-12 dataset “aat” (anaerobic-aerobic transition) (not in PRECISE-1K, but in public K-12 metadata) which took 6 time-point samples of *E. coli* from 0 to 10 minutes after aeration of a previously anaerobic chemostat culture. **A)** Top 10 iModulons by maximum difference between within-aat and PRECISE-1K activity (z-scored). For example, z-score of 5 for “Microaerobic” iModulon indicates that the maximum activity of this iModulon amongst aat samples was 5 standard deviations from the mean activity of this iModulon in PRECISE-1K. **B)** Histogram of iModulon activity across all PRECISE-1K samples and in new aat project (ArcA as example). **C)** Differential iModulon activity (DIMA) plot comparing iModulon activities at aeration onset and 10 minutes after aeration. iModulons with significant activity differences between the two time points are in blue and labeled (see **Methods** for DIMA details). **D)** iModulon activity by time from aeration (Fnr-2 as example). **E)** “Fear vs. greed” tradeoff comparing activities of RpoS (fear) and Translation (Greed) iModulons for all PRECISE-1K samples (gray) and aat samples (colored; legend in panel **F**). **F)** Tradeoff comparing activities of Fur iModulons for all PRECISE-1 K samples (gray) and aat samples (colored). Black dots indicate PRECISE-1K samples with *fur* knocked out. **G)** Tradeoff comparing activities of Fnr-2 and ArcA iModulons, with anaerobic growth conditions from PRECISE-1K in black. aat color scheme same as **F**. **H)** Principal component plot of PRECISE-1K (gray) and aat (colored; same as **F**) samples. Principal component analysis performed on iModulon activity matrix condition-wise.

Activity phase planes are another key tool for analyzing new data. The “fear vs greed” phase plane, first introduced in (Sastry et al., 2019), enables assessment of the balance of growth opportunities (underpinned by the Translation iModulon) and stresses (represented by global stress sigma factor RpoS [σ^38^]) felt by the cell in a given sample. For AAT, the aerobic transition appears to facilitate a modest increase in both fear and greed (**Figure 5E**).

The dynamic transcriptomic changes in the AAT project are more pronounced in the Fur-1/Fur-2 (**Figure 5F**) and Fnr/ArcA (**Figure 5G**) tradeoffs. As aerobic metabolism takes over, iron-related genes repressed by Fur increase in activity, as evidenced by movement of AAT samples along the Fur-1/Fur-2 axis towards the extreme activity point indicated by *fur* deletion samples. Iron is an essential component of iron-sulfur clusters that feature in many aerobic metabolism enzymes. Similarly, the Fnr/ArcA tradeoff captures the anaerobic/aerobic transition very directly. Activity of global anaerobic regulator Fnr decreases as aerobic regulator ArcA’s activity increases, with both arriving near the activity levels of PRECISE-1K’s aerobic growth control condition 10 minutes after aeration. Another viewpoint for this transition comes from a principal component analysis of PRECISE-1K and AAT conditions based on iModulon activities. Again, as time from aeration increases, samples move downward in principal component 2 (which has high weights for anaerobic metabolism iModulons) (**Figure 5H**).

Taken together, these observations highlight the essential systems-level changes occurring during aerobic transition while exemplifying PRECISE-1K’s function as an analysis resource. Further, they show the deep interpretation of TRN functions achieved through the use of iModulon activity phase planes.

## Discussion

In this work, we establish PRECISE-1K, a large, high-fidelity *E. coli* RNA-seq compendium that enables top-down transcriptional regulatory network discovery and analysis. PRECISE-1K delivers insights into the regulatory dynamics of *E. coli* at multiple scales. First, we find that gene expression levels and variance across the dataset are differentiated by factors such as core genome membership and gene function. We then present 117 regulatory modules (iModulons) that explain 56% of the total variance in the compendium and reconstitute 86% of known regulatory interactions. Thus, PRECISE-1K and its iModulons constitute the most complete top-down, computational TRN reconstruction yet generated for a microorganism. iModulons derived from PRECISE-1K cover the full range of cellular processes, from sensory two-component systems to core metabolic pathways to translation to stress responses. Thus, we demonstrate the stability, scalability, and completeness of this method for regulatory network characterization.

PRECISE-1K retains nearly all of the regulatory iModulons extracted from its predecessor PRECISE. Thus, iModulons capture fundamental regulatory modes, not dataset-specific artifacts. Increasing the dataset size nearly four-fold does not hinder regulatory network discovery; in fact, we have more than doubled the number of discovered regulatory iModulons. Taken together, PRECISE-1K and iModulons extracted from it highlight the central role that top-down, data-driven methods must take in transcriptional regulatory network discovery across organisms. Indeed, iModulons have already successfully generated top-down regulatory networks for other organisms (Chauhan et al., 2021; Lim et al., 2022; Poudel et al., 2020; Rajput et al., 2022; Rychel et al., 2020; Sastry et al., 2019; Yoo et al., 2022). The success of PRECISE-1K serves to further cement both the importance of pursuing such efforts and the reliability of the results.

Further expansion of dataset size beyond PRECISE-1K yields diminishing returns. When we added all high-quality public K-12 data to PRECISE-1K, the iModulon structure remained quite similar, with the K-12 dataset’s 124 regulatory iModulons accounting for 88% of known TRN interactions. This result highlights a key value of PRECISE-1K: this single-laboratory dataset has sufficient scale to identify stable regulatory components with robust coverage of the TRN while avoiding noise introduced by combination of data from multiple sources. As a result, we recommend usage of PRECISE-1K itself for analysis of new RNA-seq data, despite the K-12 dataset’s significantly larger scale. The lack of increase in regulatory coverage with the K-12 dataset likely reflects the need to select growth conditions that activate niche transcriptional regulators with small regulons, rather than a limitation of the top-down TRN inference method itself.

Beyond their ability to systematically characterize a TRN, iModulons have a key characteristic that bottom-up regulons lack: activity levels. This quantitative aspect of iModulons enables analysis of the functional transcriptome under specific environmental or genetic conditions. We demonstrate this capability by capturing two different functional regulatory modes of the Crp regulon based on binding site location. Differential iModulon activity (DIMA) analysis also greatly simplifies differential expression analysis; with an average of nearly twenty times fewer significantly differential variables to analyze, DIMA analysis empowers systems-level analysis of transcriptomic changes, as demonstrated in the AAT case study.

Critically, PRECISE-1K and iModulon activities enable us to discover and partially characterize putative regulons for predicted transcription factors. We demonstrate this capability by assigning a putative function in ethanol stress tolerance related to nucleotide metabolism to the YgeV regulon, based on the YgeV iModulon activation pattern. In particular, this activation coincides with knockouts of two-component system response regulators BaeR and CpxR; thus, YgeV’s role in nucleotide metabolism upon ethanol stress response may arise as a compensatory mechanism following inactivation of these more prominent TCS regulators. The specificity of this activating condition may play a role in explaining why the functions of this regulator and the genes in its regulon remain unknown. Indeed, iModulons have already proven useful in studies to characterize regulators and their regulons (Rodionova et al., 2021). PRECISE-1K likely contains other instances of untapped insights into specific regulons or regulatory dynamics, and should continue to be mined for such discoveries.

PRECISE-1K and iModulons inform experimental design beyond regulon discovery. For example, proteomics data acquisition remains more cost, labor, and time intensive than transcriptomics. Thus, it is key to identify parsimonious sets of growth conditions likely to produce the most variation in the proteome. iModulon activities can highlight growth conditions that differentially activate systems of interest, enabling judicious selection. Our knowledge base provides a centralized reference for assessment of gene expression across conditions, empowering study designs intended to perturb, modify, or delete genes.

Our example analysis of the AAT project from the K-12 dataset demonstrates perhaps the most exciting application of PRECISE-1K: analysis and contextualization of new RNA-seq datasets. PRECISE-1K’s iModulons clearly capture and summarize the regulatory dynamics at play during aerobic metabolism transition. We provide a variety of tools, both here and in our previously published code package (Sastry et al., 2021b) that will easily facilitate similar analyses for any other dataset. In this way, PRECISE-1K is not just useful in and of itself but as a backdrop for deriving regulatory insight from new data. Our example workflow for analyzing new data with PRECISE-1K - along with all other analyses from this paper - is available for use at https://github.com/SBRG/precise1k. These analyses have already enriched multi-omic studies of the aerobic respiration system (Anand et al., 2022), the adaptation of different *E. coli* strains (Kavvas et al., 2022), and the response of *E. coli* to antibiotics (Sastry et al., 2020).

Overall, PRECISE-1K and iModulons represent a critical resource for studying the transcriptional regulatory network of *E. coli.* We believe PRECISE-1K should be a standard tool for systems-level analysis of *E. coli* RNA-seq data from all sources. As the number of publicly available datasets increases for other microorganisms, this study serves as a roadmap for interrogating similar datasets for less characterized organisms, with the potential to yield equally impactful insights into those organisms’ regulatory network structures.

## Methods

### RNA-seq processing and quality control

PRECISE-1K consists of all data in PRECISE (Sastry et al., 2019), along with additional data generated in the Palsson Lab, some previously published and some published for the first time here.

Starting from 1055 candidate samples, data was processed using a Nextflow (Di Tommaso et al., 2017) [pipeline](https://github.com/avsastry/modulome-workflow) designed for processing microbial RNA-seq datasets (Sastry et al., 2021b), and run on Amazon Web Services (AWS) Batch.

First, raw read trimming was performed using [Trim Galore](https://www.bioinformatics.babraham.ac.uk/projects/trim_galore/) with the default options, followed by [FastQC](http://www.bioinformatics.babraham.ac.uk/projects/fastqc/) on the trimmed reads. Next, reads were aligned to the *E. coli* K-12 MG1655 reference genome (RefSeq accession number NC_000913.3) using Bowtie (Langmead et al., 2009). The read direction was inferred using RSEQC (Wang et al., 2012) before generating read counts using featureCounts (Liao et al., 2014). Finally, all quality control metrics were compiled using MultiQC (Ewels et al., 2016) and the final expression dataset was reported in units of log-transformed Transcripts Per Million (log-TPM).

Samples were considered “high-quality” if they met all of the following criteria:

- “Pass” on the all of the following FastQC checks: *per_base_sequence_quality, per_sequence_quality_scores, per_base_n_content, adapter_content*
- At least 500,000 reads mapped to coding sequences (CDS) from the reference genome (NC_000913.3)
- Not an outlier in hierarchical clustering based on pairwise Pearson correlation between all samples (outlier defined as cluster with number of samples <1% of the total number of samples)
- Minimum Pearson correlation with biological replicates (if any) 0.95 (if more than 2 biological replicates, keep samples with high correlation in “greedy” manner, dropping samples that have at least one sub-threshold correlation with all other replicates)

Following this processing and QC workflow, 1035 high-quality RNA-seq samples remained. These samples and their metadata define PRECISE-1K. log-TPM, raw read count, QC data files, and sample metadata for all 1055 original samples are included in the supplementary data. These files may also be found in the data directory of this project’s GitHub repository.

### Differentially expressed gene (DEG) computation

Differentially expressed genes (DEGs) were identified using the *DESeq2* package (Love et al., 2014) on the PRECISE-1K RNA-seq dataset. Genes with a log_2_ fold change greater than 1.5 and a false discovery rate (FDR) value less than 0.05 were considered to be differentially expressed genes. Genes with p-values assigned “NA” based on extreme count outlier detection were not considered as potential DEGs. The number of DEGs was computed for each unique pair of conditions within each project in PRECISE-1K, for a total of 6104 pairwise computations.

### iModulon computation

To compute the optimal set of independent components (iModulons), the previously described OptICA method (McConn et al., 2021) was run on PRECISE-1K. The iModulon matrices **M** and **A** (along with iModulon metadata containing annotation as described below) are available in the supplementary data files and in this project’s GitHub repository.

### Differential iModulon activity (DIMA) computation

Differentially activated iModulons were computed with a similar process as previously detailed (Sastry et al., 2019). For each iModulon, the average activity of the iModulon between biological replicates, if available, was computed. Then, the absolute value of the difference in iModulon activities between the two conditions was compared to the fitted log-normal distribution of all differences in activity for the iModulon. iModulons that had an absolute value of activity greater than 5, and an FDR below 0.05 were considered to be significant. The number of DIMAs was computed for each unique pair of conditions within each project in the PRECISE-1K compendium, mirroring DEG computation.

### iModulon annotation

First, a gold-standard TRN reference annotation was downloaded from RegulonDB v10.5 (Santos-Zavaleta et al., 2019). For each iModulon, enrichment of the set of genes in the iModulon against known RegulonDB regulons was computed using Fisher’s Exact Test, with false discovery rate controlled at 10^-5^ using the Benjamini-Hochberg correction. By default, iModulons were compared to all possible single regulons and all possible combinations of two regulons (intersection only). The regulons used by default consisted of only strong and confirmed evidence regulatory interactions, per RegulonDB. When multiple significant enrichments were available, the enrichment with the lowest adjusted *P* value was used for annotation. In the event of near equal *P* values (within an order of magnitude) across multiple enrichments, the priority was given to intersection regulons, followed by single regulons, followed by union regulons. If no significant enrichments were available, the following adjustments were used, in this order: relax evidence requirement to include weak evidence regulatory interactions; search only for single regulon enrichments; allow up to 3 regulons to be combined for enrichment; allow regulon unions as well as intersections (with priority given to intersections). If the iModulon consisted of genes with annotated co-regulation by 4 or more genes, a specific enrichment calculation was made to determine the enrichment statistics. If none of these adjustments yielded a significant enrichment, the iModulon was annotated as non-regulatory. All parameters and statistics related to calculation of TRN enrichments for regulatory iModulons are recorded in the iModulon metadata table, available in the GitHub repository.

iModulons were named and annotated according to the following ruleset:

<u>General</u>

- Rule #1: iModulon names must be fewer than ~15 characters
- Rule #2: iModulon names must be unique. If iModulons would otherwise have the same name, append “-1”, “-2”, etc., as needed to disambiguate. By default, order the suffixes by decreasing explained variance, unless another numbering is specifically preferred (e.g. aligning Crp-1 and Crp-2 with Crp binding site classes).

<u>Case 1 - Regulatory</u>

The iModulon has a significant regulon enrichment chosen as described above:

- Rule #1: Name the iModulon after the primary function of the associated regulon(s) (e.g. the iModulon enriched for the CdaR regulon is named “Sugar Diacid”)
- Rule #2: If no clear primary function is available for the iModulon, name the iModulon directly after the enriched regulon (e.g. the iModulon enriched for the CpxR regulon is named “CpxR”, as CpxR controls a diverse set of functions).
- Exception #1: if the enriched regulon corresponds to a well-known global regulator (i.e. Fur, CRP, RpoS), name the iModulon after that regulator.
- Exception #2: if the name per Rule #1 would violate General Rule #1, name the iModulon directly after the enriched regulon (e.g. the iModulon enriched for the union of the FucR and ExuR regulons is named “FucR/ExuR” instead of “Fucose/Galacturonate/Glucuronate”)
- Exception #3: if applying Rule #2, and the regulon enrichment involves an intersection between a global regulator and a local regulator (i.e. cooperative regulation), the global regulator may be dropped from the name (e.g. “NtrC-1” instead of “RpoN+NtrC-1”, as RpoN is a larger-regulon sigma factor which co-regulates with the more-specific NtrC).

<u>Case 2 - Genomic</u>

The iModulon activity profile has a clear correlation with a sample involving a specific genetic or genomic intervention:

- Rule #1: if the iModulon captures intentional knockout of a gene (e.g. *geneA* is knocked out in *sampleA,* and the iModulon has a large positive gene weight for *geneA* and a large negative activity level for *sampleA,* accounting for the lack of *geneA* expression in *sampleA),* name the iModulon “[gene name] KO” (e.g. baeR KO)
- Rule #2: Similarly, if the iModulon captures intentional overexpression of a particular gene, name the iModulon “[gene name] OE” (e.g. “malE OE”)
- Rule #3: if the iModulon captures expression changes in relation to evolved samples (ALE), as determined by comparing the iModulon activities to known ALE samples, name the iModulon “[name of ALE project] Del” (for deletions), “[name of ALE project] Amp” (for amplifications), or “name of ALE project] Mut” (for mixed effect mutations) (e.g. ROS TALE Del-1)
- Rule #4: if the iModulon also has a significant regulon enrichment as described above, prioritize the specific genetic/genomic change.

<u>Case 3 - Biological</u>

The iModulon does not have a significant regulon enrichment, does not relate to a specific genetic or genomic change, but the member genes share a clear biological function:

- Rule #1: Name the iModulon after the shared biological function (e.g. the “LPS” iModulon consists of many genes related to lipopolysaccharide biosynthesis and export, though no significant regulon enrichment was found for this iModulon’s genes).

<u>Case 4 - Single-Gene Dominant</u>

The iModulon contains one specific gene with a gene weight at least twice as large as the next closest gene, does not fall into Case 2 - Genomic, and contains only the one highly-weighted genes, or at most 5 other genes with gene weights very close to the iModulon’s threshold

- Rule #1: Name the iModulon after the dominant gene (e.g. the “ymdG” iModulon consists solely of the *ymdG* gene)

<u>Case 5 - Uncharacterized</u>

The iModulon does not meet any of the previous criteria for naming

- Rule #1: Name the iModulon “UC-#” (short for “Uncharacterized”), with the number incrementing for each uncharacterized iModulon.

### Compiling the K-12 public dataset

Data was compiled from NCBI SRA as described previously (Sastry et al., 2021b). Initially, all data annotated as RNA-seq for *E. coli* was inspected. RNA-seq samples were discarded if the strain was not from a K-12 strain, if the strain was missing, or if the type of experiment was not actually RNA-seq. After initial curation, data was processed and quality controlled as described previously, and iModulons were computed in the same manner as described above.

## Data and Code Availability

All data and codes are available at https://github.com/SBRG/precise1k.

## Supplemental Figures

**Supplemental Figure 1:**
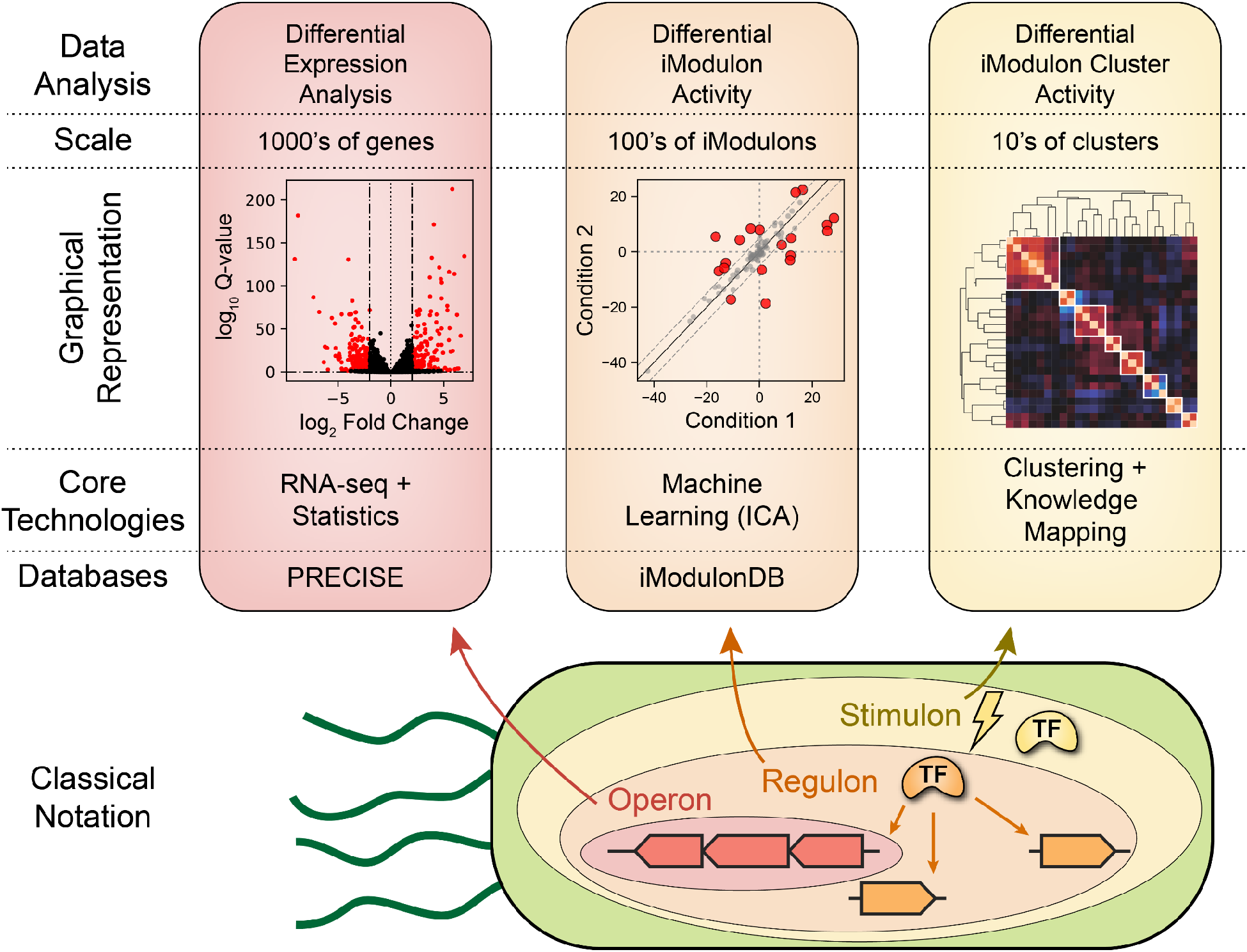
Multi-scale analysis of PRECISE-1K. The levels of analysis approximately correspond to the definition of an operon and a regulon, and also a quantitative definition of the notion of a stimulon. The ‘scale’ indicates the reduction of dimensionality over the levels shown.

**Supplemental Figure 2:**
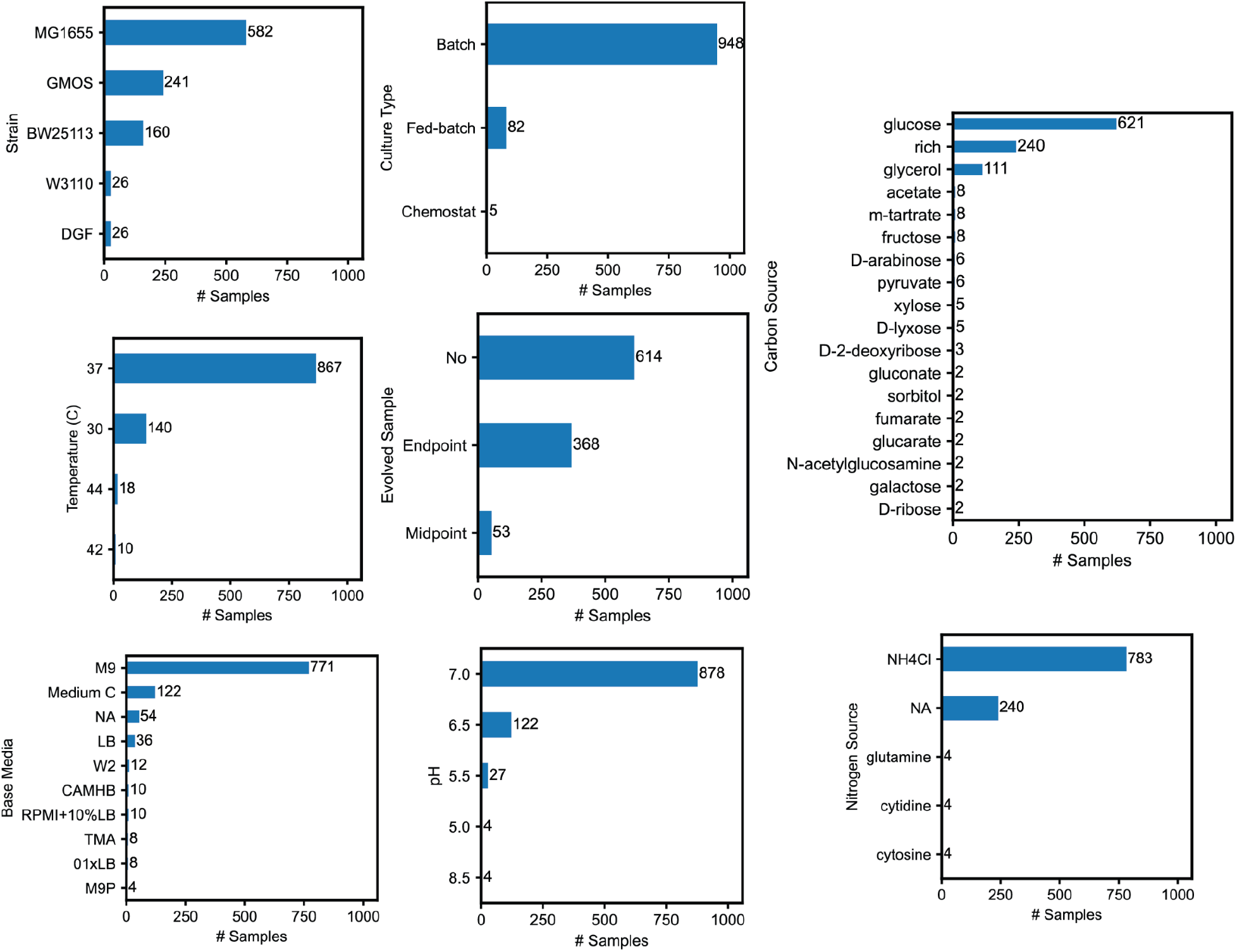
Breakdown of major growth conditions for PRECISE-1K.

**Supplemental Figure 3:**
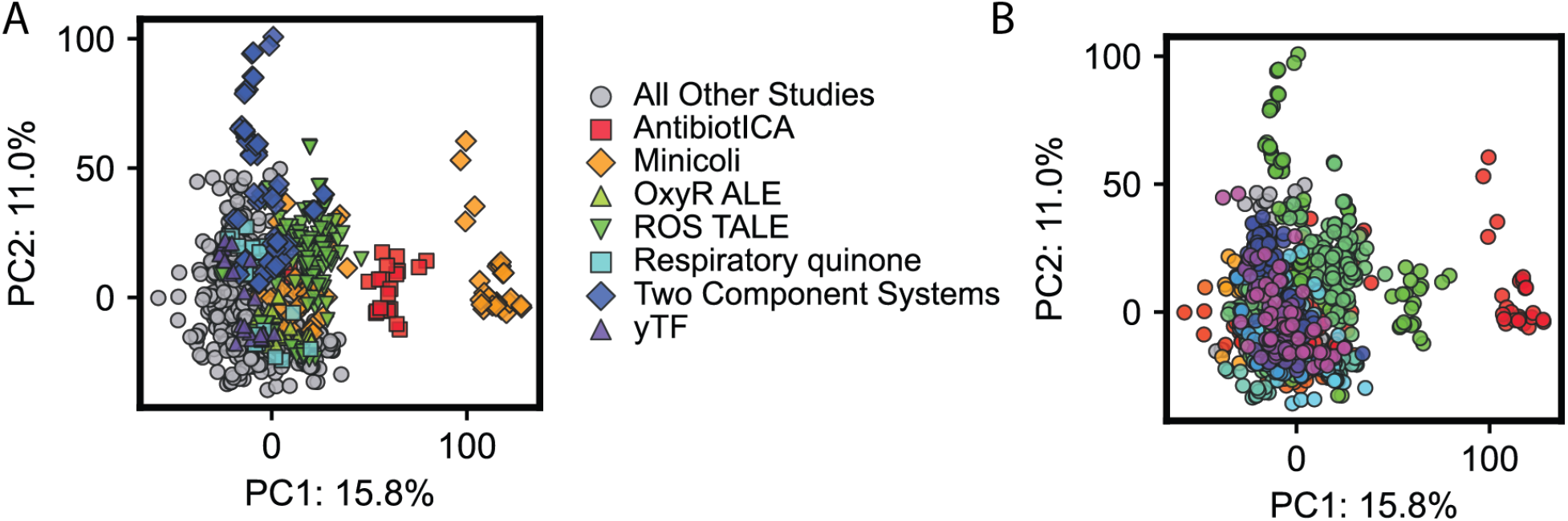
Principal component analysis (PCA) of PRECISE-1K. **A)** First 2 principal components, colored by project (n=1035 samples). **B)** First 2 principal components, colored by each of 21 distinct RNA-seq library preparers.

**Supplemental Figure 4:**
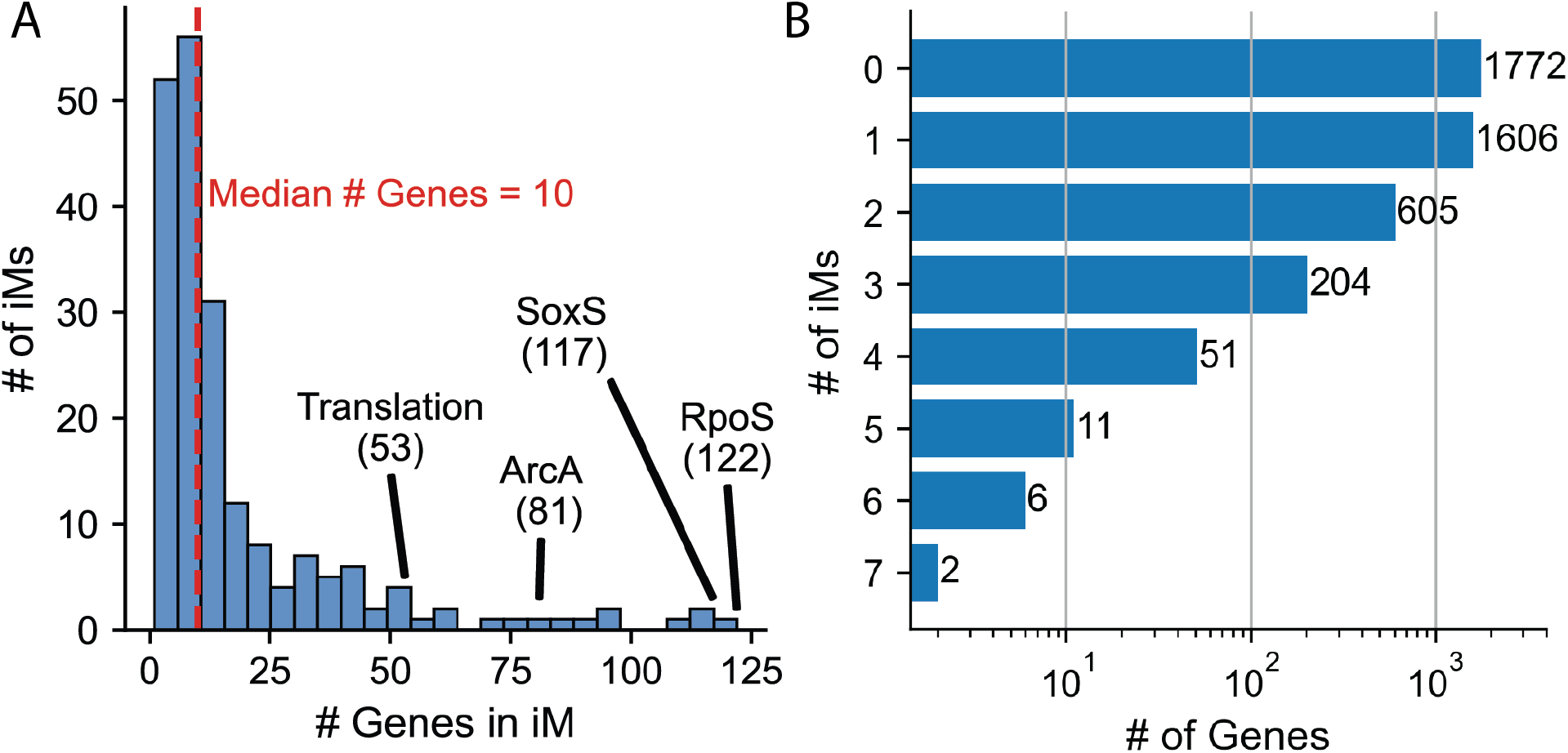
iModulon gene membership breakdown. **A)** Histogram of iModulon sizes. n=201 iModulons. **B)** Breakdown of genes by number of iModulons of which they are a member. n=4257 genes. 2485 genes are members of at least 1 iModulon.

**Supplemental Figure 5:**
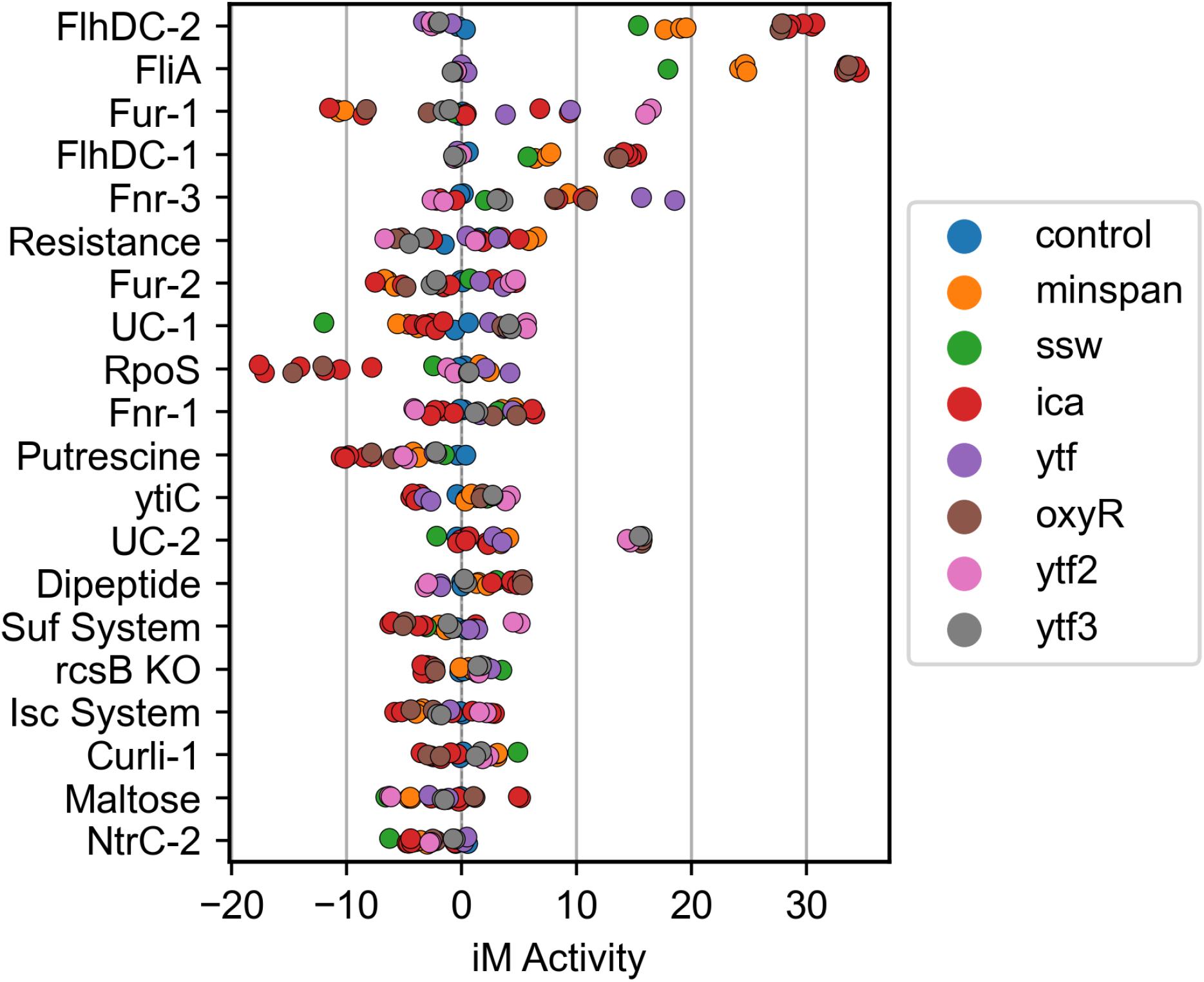
Most variant iModulon activities in control conditions across projects. iModulon activities with the highest median absolute deviation across 20 samples of wild-type growth in M9 medium with glucose across 8 projects are displayed.

**Supplemental Figure 6:**
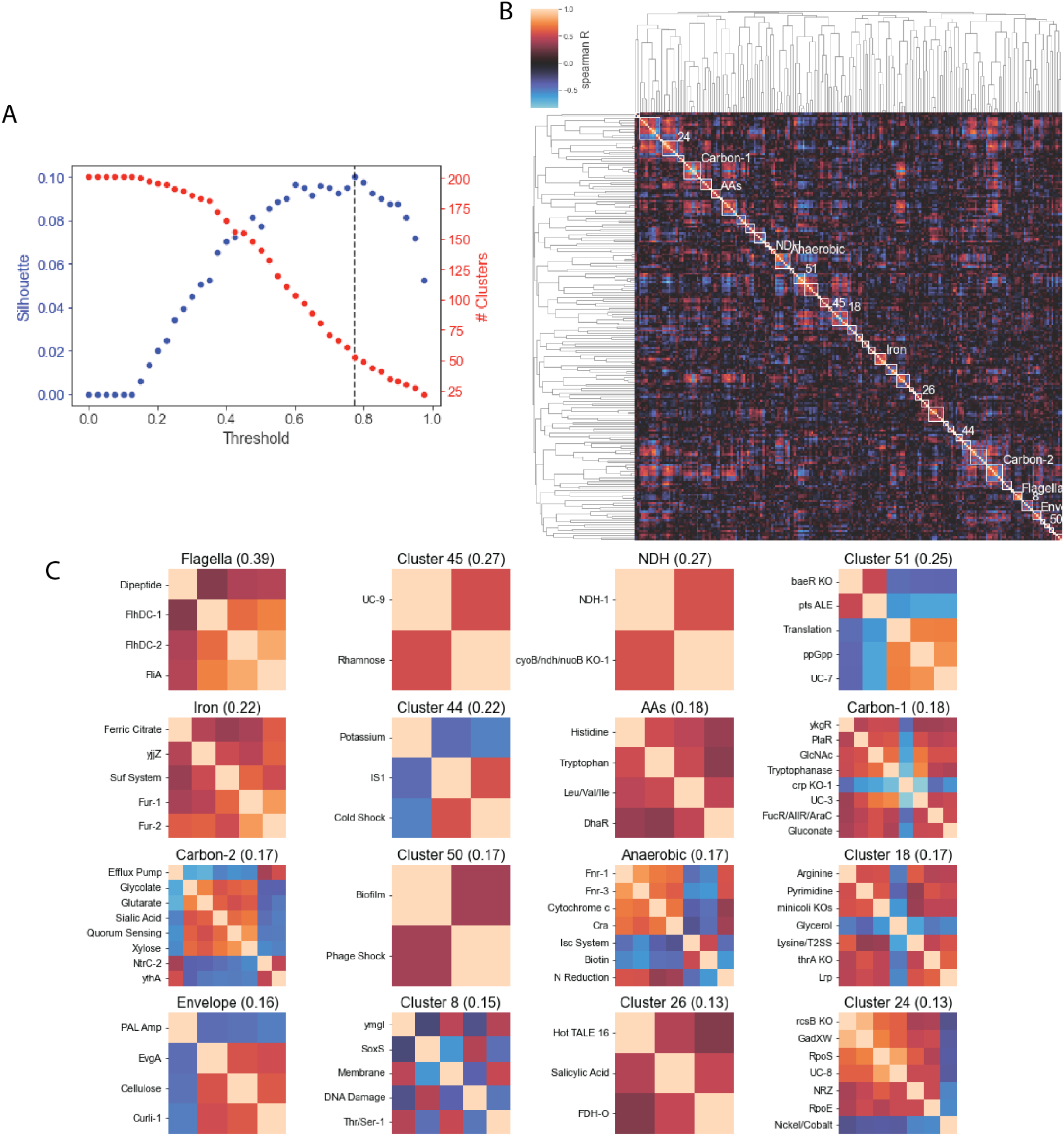
iModulon activity clustering for PRECISE-1K: defining stimulons. **(A)** Automatic distance threshold determination. Distance thresholds for determination of optimal clusters after hierarchical clustering were determined by scanning possible distance thresholds and computing a metric of cluster distinctness - the silhouette score - at each possible threshold. Black dashed line indicates final choice of distance threshold (0.725). **(B)** Clustermap of PRECISE-1K iModulon activities. Colorbar indicates Spearman’s correlation coefficient, the distance metric used to perform pairwise iModulon activity comparisons. 16 most distinct clusters - by silhouette score - are labeled with yellow text corresponding to panel **C**. **(C)** 16 most distinct clusters of iModulon activities from PRECISE-1K (silhouette scores parenthesized).

**Supplemental Figure 7:**
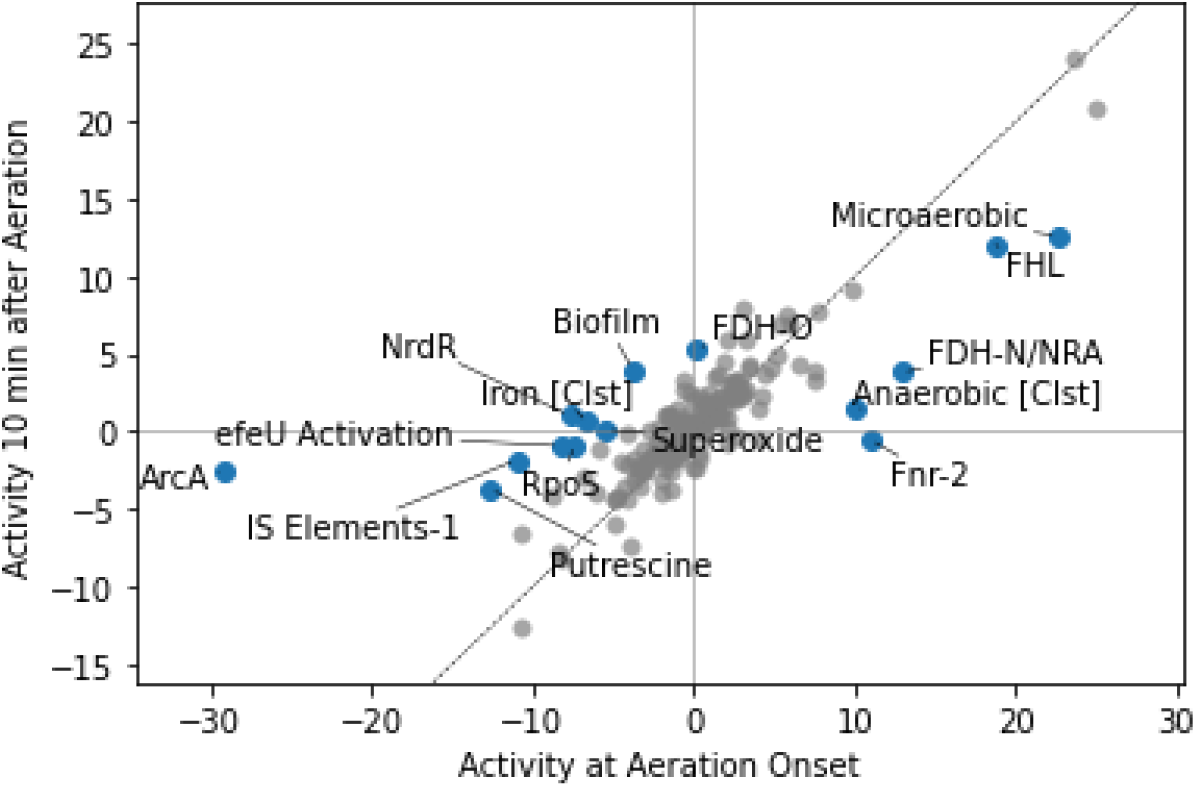
Differential iModulon activity (DIMA) plot between onset of aeration and 10 minutes post-aeration with activity clusters (stimulons) included (indicated with [Clst] suffix).

## Notes

https://github.com/SBRG/precise1k

### Competing Interest Statement

The authors have declared no competing interest.

### Summary of Updates

This manuscript has been modified to reflect an updated/expanded version of the underlying dataset. The analyses in the manuscript have been repeated on this larger dataset. Certain figures have been modified for clarity. A section on use cases for the dataset has also been added.

## Bibliography

Anand, A., Chen, K., Yang, L., Sastry, A.V., Olson, C.A., Poudel, S., Seif, Y., Hefner, Y., Phaneuf, P.V., Xu, S., et al. (2019). Adaptive evolution reveals a tradeoff between growth rate and oxidative stress during naphthoquinone-based aerobic respiration. Proc. Natl. Acad. Sci. U. S. A. 116, 25287–25292..

Anand, A., Chen, K., Catoiu, E., Sastry, A.V., Olson, C.A., Sandberg, T.E., Seif, Y., Xu, S., Szubin, R., Yang, L., et al. (2020). OxyR Is a Convergent Target for Mutations Acquired during Adaptation to Oxidative Stress-Prone Metabolic States. Mol. Biol. Evol. 37, 660–667..

Anand, A., Patel, A., Chen, K., Olson, C.A., Phaneuf, P.V., Lamoureux, C., Hefner, Y., Szubin, R., Feist, A.M., and Palsson, B.O. (2022). Laboratory evolution of synthetic electron transport system variants reveals a larger metabolic respiratory system and its plasticity. Nat. Commun. 13, 3682..

Avsec, Ž., Agarwal, V., Visentin, D., Ledsam, J.R., Grabska-Barwinska, A., Taylor, K.R., Assael, Y., Jumper, J., Kohli, P., and Kelley, D.R. (2021). Effective gene expression prediction from sequence by integrating long-range interactions. Nat. Methods 18, 1196–1203..

Bui, T.T., and Selvarajoo, K. (2020). Attractor Concepts to Evaluate the Transcriptome-wide Dynamics Guiding Anaerobic to Aerobic State Transition in Escherichia coli. Sci. Rep. 10, 5878.

Busby, S., and Ebright, R.H. (1999). Transcription activation by catabolite activator protein (CAP). J. Mol. Biol. 293, 199–213..

Chauhan, S.M., Poudel, S., Rychel, K., Lamoureux, C., Yoo, R., Al Bulushi, T., Yuan, Y., Palsson, B.O., and Sastry, A.V. (2021). Machine Learning Uncovers a Data-Driven Transcriptional Regulatory Network for the Crenarchaeal Thermoacidophile Sulfolobus acidocaldarius. Front. Microbiol. 12, 753521..

Chen, K., Anand, A., Olson, C., Sandberg, T.E., Gao, Y., Mih, N., and Palsson, B.O. (2021). Bacterial fitness landscapes stratify based on proteome allocation associated with discrete aero-types. PLoS Comput. Biol. 17, e1008596..

Choudhary, K.S., Kleinmanns, J.A., Decker, K., Sastry, A.V., Gao, Y., Szubin, R., Seif, Y., and Palsson, B.O. (2020). Elucidation of Regulatory Modes for Five Two-Component Systems in Escherichia coli Reveals Novel Relationships. mSystems 5. https://doi.org/10.1128/mSystems.00980-20.

Comon, P. (1994). Independent component analysis, A new concept? Signal Processing 36, 287–314..

DeLisa, M.P., Wu, C.F., Wang, L., Valdes, J.J., and Bentley, W.E. (2001). DNA microarray-based identification of genes controlled by autoinducer 2-stimulated quorum sensing in Escherichia coli. J. Bacteriol. 183, 5239–5247..

Di Tommaso, P., Chatzou, M., Floden, E.W., Barja, P.P., Palumbo, E., and Notredame, C. (2017). Nextflow enables reproducible computational workflows. Nat. Biotechnol. 35, 316–319..

Du, B., Olson, C.A., Sastry, A.V., Fang, X., Phaneuf, P.V., Chen, K., Wu, M., Szubin, R., Xu, S., Gao, Y, et al. (2020). Adaptive laboratory evolution of Escherichia coli under acid stress. Microbiology 166, 141–148..

ENCODE Project Consortium (2012). An integrated encyclopedia of DNA elements in the human genome. Nature 489, 57–74..

Ewels, P., Magnusson, M., Lundin, S., and Käller, M. (2016). MultiQC: summarize analysis results for multiple tools and samples in a single report. Bioinformatics 32, 3047–3048..

Gao, Y., Yurkovich, J.T., Seo, S.W., Kabimoldayev, I., Dräger, A., Chen, K., Sastry, A.V., Fang, X., Mih, N., Yang, L., et al. (2018). Systematic discovery of uncharacterized transcription factors in Escherichia coli K-12 MG1655. Nucleic Acids Res. https://doi.org/10.1093/nar/gky752.

Ghatak, S., King, Z.A., Sastry, A., and Palsson, B.O. (2019). The y-ome defines the 35% of Escherichia coli genes that lack experimental evidence of function. Nucleic Acids Res. 47, 2446–2454..

GTEx Consortium (2015). Human genomics. The Genotype-Tissue Expression (GTEx) pilot analysis: multitissue gene regulation in humans. Science 348, 648–660..

Heckmann, D., Lloyd, C.J., Mih, N., Ha, Y., Zielinski, D.C., Haiman, Z.B., Desouki, A.A., Lercher, M.J., and Palsson, B.O. (2018). Machine learning applied to enzyme turnover numbers reveals protein structural correlates and improves metabolic models. Nat. Commun. 9. https://doi.org/10.1038/s41467-018-07652-6.

Heckmann, D., Campeau, A., Lloyd, C.J., Phaneuf, P.V., Hefner, Y., Carrillo-Terrazas, M., Feist, A.M., Gonzalez, D.J., and Palsson, B.O. (2020). Kinetic profiling of metabolic specialists demonstrates stability and consistency of in vivo enzyme turnover numbers. Proc. Natl. Acad. Sci. U. S. A. 117, 23182–23190..

Hirokawa, Y., Kawano, H., Tanaka-Masuda, K., Nakamura, N., Nakagawa, A., Ito, M., Mori, H., Oshima, T., and Ogasawara, N. (2013). Genetic manipulations restored the growth fitness of reduced-genome Escherichia coli. J. Biosci. Bioeng. 116, 52–58..

Kavvas, E.S., Long, C.P., Sastry, A., Poudel, S., Antoniewicz, M.R., Ding, Y., Mohamed, E.T., Szubin, R., Monk, J.M., Feist, A.M., et al. (2022). Experimental Evolution Reveals Unifying Systems-Level Adaptations but Diversity in Driving Genotypes. mSystems e0016522..

Kelley, D.R., Reshef, Y.A., Bileschi, M., Belanger, D., McLean, C.Y., and Snoek, J. (2018). Sequential regulatory activity prediction across chromosomes with convolutional neural networks. Genome Res. 28, 739–750..

Kwon, M.S., Lee, B.T., Lee, S.Y., and Kim, H.U. (2020). Modeling regulatory networks using machine learning for systems metabolic engineering. Curr. Opin. Biotechnol. 65, 163–170..

Langmead, B., Trapnell, C., Pop, M., and Salzberg, S.L. (2009). Ultrafast and memory-efficient alignment of short DNA sequences to the human genome. Genome Biol. 10, R25..

Lawson, C.L., Swigon, D., Murakami, K.S., Darst, S.A., Berman, H.M., and Ebright, R.H. (2004). Catabolite activator protein: DNA binding and transcription activation. Curr. Opin. Struct. Biol. 14, 10–20..

Leader, D.P., Krause, S.A., Pandit, A., Davies, S.A., and Dow, J.A.T. (2018). FlyAtlas 2: a new version of the Drosophila melanogaster expression atlas with RNA-Seq, miRNA-Seq and sex-specific data. Nucleic Acids Res. 46, D809–D815..

Leinonen, R., Sugawara, H., Shumway, M., and International Nucleotide Sequence Database Collaboration (2011). The sequence read archive. Nucleic Acids Res. 39, D19–D21..

Liao, Y., Smyth, G.K., and Shi, W. (2014). featureCounts: an efficient general purpose program for assigning sequence reads to genomic features. Bioinformatics 30, 923–930..

Lim, H.G., Rychel, K., Sastry, A.V., Bentley, G.J., Mueller, J., Schindel, H.S., Larsen, P.E., Laible, P.D., Guss, A.M., Niu, W., et al. (2022). Machine-learning from Pseudomonas putida KT2440 transcriptomes reveals its transcriptional regulatory network. Metab. Eng. 72, 297–310.

Liu, and Markatou Evaluation of methods in removing batch effects on RNA-seq data. Infect. Dis. Transl. Med.

Love, M.I., Huber, W., and Anders, S. (2014). Moderated estimation of fold change and dispersion for RNA-seq data with DESeq2. Genome Biol. 15, 550..

McCloskey, D., Xu, S., Sandberg, T.E., Brunk, E., Hefner, Y., Szubin, R., Feist, A.M., and Palsson, B.O. (2018). Evolution of gene knockout strains of E. coli reveal regulatory architectures governed by metabolism. Nat. Commun. 9, 3796..

McConn, J.L., Lamoureux, C.R., Poudel, S., Palsson, B.O., and Sastry, A.V. (2021). Optimal dimensionality selection for independent component analysis of transcriptomic data.

Mehta, P., Casjens, S., and Krishnaswamy, S. (2004). Analysis of the lambdoid prophage element e14 in the E. coli K-12 genome. BMC Microbiol. 4, 4..

Norsigian, C.J., Pusarla, N., McConn, J.L., Yurkovich, J.T., Dräger, A., Palsson, B.O., and King, Z. (2020). BiGG Models 2020: multi-strain genome-scale models and expansion across the phylogenetic tree. Nucleic Acids Res. 48, D402–D406..

Poudel, S., Tsunemoto, H., Seif, Y., Sastry, A.V., Szubin, R., Xu, S., Machado, H., Olson, C.A., Anand, A., Pogliano, J., et al. (2020). Revealing 29 sets of independently modulated genes in Staphylococcus aureus, their regulators, and role in key physiological response. Proc. Natl. Acad. Sci. U. S. A. 117, 17228–17239..

Qiu, S., Lamoureux, C., Akbari, A., Palsson, B.O., and Zielinski, D.C. (2022). Quantitative sequence basis for the E. coli transcriptional regulatory network.

Rajput, A., Tsunemoto, H., Sastry, A.V., Szubin, R., Rychel, K., Sugie, J., Pogliano, J., and Palsson, B.O. (2022). Machine learning from Pseudomonas aeruginosa transcriptomes identifies independently modulated sets of genes associated with known transcriptional regulators. Nucleic Acids Res. 50, 3658–3672..

Reitzer, L., and Schneider, B.L. (2001). Metabolic Context and Possible Physiological Themes of ς54-Dependent Genes in Escherichia coli. Microbiol. Mol. Biol. Rev. 65, 422–444..

Rodionova, I.A., Gao, Y., Sastry, A., Yoo, R., Rodionov, D.A., Saier, M.H., and Palsson, B.Ø. (2020a). Synthesis of the novel transporter YdhC, is regulated by the YdhB transcription factor controlling adenosine and adenine uptake.

Rodionova, I.A., Gao, Y., Sastry, A.V., Monk, J., and Szubin, R. 2020b). PtrR (YneJ) is a novel E. coli transcription factor regulating the putrescine stress response and glutamate utilization. bioRxiv.

Rodionova, I.A., Gao, Y., Sastry, A., Hefner, Y., Lim, H.G., Rodionov, D.A., Saier, M.H., Jr, and Palsson, B.O. (2021). Identification of a transcription factor, PunR, that regulates the purine and purine nucleoside transporter punC in E. coli. Commun Biol 4, 991..

Rychel, K., Sastry, A.V., and Palsson, B.O. (2020). Machine learning uncovers independently regulated modules in the Bacillus subtilis transcriptome. Nat. Commun. 11, 6338..

Rychel, K., Decker, K., Sastry, A.V., Phaneuf, P.V., Poudel, S., and Palsson, B.O. (2021). iModulonDB: a knowledgebase of microbial transcriptional regulation derived from machine learning. Nucleic Acids Res. 49, D112–D120..

Saelens, W., Cannoodt, R., and Saeys, Y. (2018). A comprehensive evaluation of module detection methods for gene expression data. Nat. Commun. 9. https://doi.org/10.1038/s41467-018-03424-4.

Sandberg, T.E., Szubin, R., Phaneuf, P.V., and Palsson, B.O. (2020). Synthetic cross-phyla gene replacement and evolutionary assimilation of major enzymes. Nat Ecol Evol 4, 1402–1409.

Santos-Zavaleta, A., Salgado, H., Gama-Castro, S., Sánchez-Pérez, M., Gómez-Romero, L., Ledezma-Tejeida, D., García-Sotelo, J.S., Alquicira-Hernández, K., Muñiz-Rascado, L.J., Peña-Loredo, P., et al. (2019). RegulonDB v 10.5: Tackling challenges to unify classic and high throughput knowledge of gene regulation in E. coli K-12. Nucleic Acids Res. 47, D212–D220..

Sastry, A., Dillon, N., Poudel, S., Hefner, Y., Xu, S., Szubin, R., Feist, A., Nizet, V., and Palsson, B. (2020). Decomposition of transcriptional responses provides insights into differential antibiotic susceptibility.

Sastry, A.V., Gao, Y., Szubin, R., Hefner, Y., Xu, S., Kim, D., Choudhary, K.S., Yang, L., King, Z.A., and Palsson, B.O. (2019). The Escherichia coli transcriptome mostly consists of independently regulated modules. Nat. Commun. 10, 5536..

Sastry, A.V., Hu, A., Heckmann, D., Poudel, S., Kavvas, E., and Palsson, B.O. (2021a). Independent component analysis recovers consistent regulatory signals from disparate datasets. PLoS Comput. Biol. 17, e1008647..

Sastry, A.V., Poudel, S., Rychel, K., Yoo, R., Lamoureux, C.R., Chauhan, S., Haiman, Z.B., Al Bulushi, T., Seif, Y., and Palsson, B.O. (2021b). Mining all publicly available expression data to compute dynamic microbial transcriptional regulatory networks.

Schmidt, A., Kochanowski, K., Vedelaar, S., Ahrné, E., Volkmer, B., Callipo, L., Knoops, K., Bauer, M., Aebersold, R., and Heinemann, M. (2016). The quantitative and condition-dependent Escherichia coli proteome. Nat. Biotechnol. 34, 104–110..

Tan, J., Sastry, A.V., Fremming, K.S., Bjørn, S.P., Hoffmeyer, A., Seo, S., Voldborg, B.G., and Palsson, B.O. (2020). Independent component analysis of E. coli’s transcriptome reveals the cellular processes that respond to heterologous gene expression. Metab. Eng. 61, 360–368..

Touati, D., Jacques, M., Tardat, B., Bouchard, L., and Despied, S. (1995). Lethal oxidative damage and mutagenesis are generated by iron in delta fur mutants of Escherichia coli: protective role of superoxide dismutase. J. Bacteriol. 177, 2305–2314..

Wang, L., Wang, S., and Li, W. (2012). RSeQC: quality control of RNA-seq experiments. Bioinformatics 28, 2184–2185..

Yoo, R., Rychel, K., Poudel, S., Al-Bulushi, T., Yuan, Y., Chauhan, S., Lamoureux, C., Palsson, B.O., and Sastry, A. (2022). Machine Learning of All Mycobacterium tuberculosis H37Rv RNA-seq Data Reveals a Structured Interplay between Metabolism, Stress Response, and Infection. mSphere 7, e0003322..

Zhang, Y., Parmigiani, G., and Johnson, W.E. (2020). ComBat-seq: batch effect adjustment for RNA-seq count data. NAR Genom Bioinform 2, lqaa078..

Zhang, Z., Pan, Z., Ying, Y., Xie, Z., Adhikari, S., Phillips, J., Carstens, R.P., Black, D.L., Wu, Y., and Xing, Y. (2019). Deep-learning augmented RNA-seq analysis of transcript splicing. Nat. Methods 16, 307–310..

Ziemann, M., Kaspi, A., and El-Osta, A. (2019). Digital expression explorer 2: a repository of uniformly processed RNA sequencing data. Gigascience 8. https://doi.org/10.1093/gigascience/giz022.

Zrimec, J., Börlin, C.S., Buric, F., Muhammad, A.S., Chen, R., Siewers, V., Verendel, V., Nielsen, J., Töpel, M., and Zelezniak, A. (2020). Deep learning suggests that gene expression is encoded in all parts of a co-evolving interacting gene regulatory structure. Nat. Commun. 11, 6141..

